# Roles of Tubulin Concentration during Prometaphase and Ran-GTP during Anaphase of *C. elegans* meiosis

**DOI:** 10.1101/2024.04.19.590357

**Authors:** Ting Gong, Karen L. McNally, Siri Konanoor, Alma Peraza, Cynthia Bailey, Stefanie Redemann, Francis J. McNally

**Affiliations:** Department of Molecular and Cellular Biology University of California, Davis Davis, CA 95616 USA; Department of Cell Biology University of Virginia, School of Medicine Charlottesville, VA, USA

## Abstract

In many animal species, the oocyte meiotic spindle, which is required for chromosome segregation, forms without centrosomes. In some systems, Ran-GEF on chromatin initiates spindle assembly. We found that in *C. elegans* oocytes, endogenously-tagged Ran-GEF dissociates from chromatin during spindle assembly but re-associates during meiotic anaphase. Meiotic spindle assembly occurred after auxin-induced degradation of Ran-GEF but anaphase I was faster than controls and extrusion of the first polar body frequently failed. In search of a possible alternative pathway for spindle assembly, we found that soluble tubulin concentrates in the nuclear volume during germinal vesicle breakdown. We found that the concentration of soluble tubulin in the metaphase spindle region is enclosed by ER sheets which exclude cytoplasmic organelles including mitochondria and yolk granules. Measurement of the volume occupied by yolk granules and mitochondria indicated that volume exclusion would be sufficient to explain the concentration of tubulin in the spindle volume. We suggest that this concentration of soluble tubulin may be a redundant mechanism promoting spindle assembly near chromosomes.

## Introduction

Errors in chromosome segregation result in aneuploidy, a leading cause of embryonic lethality, and congenital defects if they occur during meiosis and cancer if they occur during mitosis (Nasmyth, 2002). Faithfull chromosome segregation in most eukaryotes relies on a bipolar spindle segregating chromosomes into daughter cells. The bipolar spindle is composed of thousands of microtubules, whose organization and stability are dynamically regulated to ensure proper chromosome attachment, alignment and segregation (Bennabi et al., 2016; Kline-Smith and Walczak, 2004; Mullen et al., 2019).

In most mitotic cells, centrosomes at the two spindle poles act as major microtubule organization centers (MTOCs), in which spindle assembly factors (SAFs) are recruited to nucleate spindle microtubules (Petry, 2016; Prosser and Pelletier, 2017). Each centrosome contains a pair of centrioles and surrounding pericentriolar material (PCM) proteins (Bornens, 2012; Hinchcliffe, 2014; Kellogg et al., 2003; Sanchez and Feldman, 2017; Wang et al., 2014). However, mitotic cells lacking centrosomes can sometimes still assemble bipolar spindles, indicating the existence of additional pathways in spindle formation (Conduit et al., 2015; Khodjakov et al., 2000; Prosser and Pelletier, 2017). Moreover, centrosomes gradually degenerate during oogenesis, and female meiotic spindles in many animal species form without centrosomes (Dumont and Desai, 2012; Gruss, 2018; Heald et al., 1997; Mikeladze-Dvali et al., 2012; Schuh and Ellenberg, 2007). It is well-known that human oocytes, especially from individuals with advanced maternal ages or in vitro fertilizations (IVF) are highly prone to meiotic spindle formation errors, resulting in aneuploid embryos (Angell, 1991; Fair and Lonergan, 2023; Thomas et al., 2021).

Two general pathways have been proposed to replace centrosomes and nucleate microtubules for spindle formation in oocytes, acentriolar cytoplasmic MTOCs and chromosome-directed spindle assembly (Li et al., 2006; Schuh and Ellenberg, 2007; Wu et al., 2022). In mouse oocytes, multiple de novo MTOCs originate from cytoplasmic microtubules prior to GVBD, which later increase in number and cluster into a multipolar spindle. These MTOCs lack centrioles but are enriched in PCM proteins (Schuh and Ellenberg, 2007). Cytoplasmic non-centrosomal MTOC-like structures enriched with different proteins have been observed in human oocytes, driven by microtubule associated protein TACC3 (Wu et al., 2022). Non-centrosomal MTOCs have not been reported in oocytes of *Drosophila* or *C. elegans*.

Chromosome-directed spindle assembly has been studied extensively in *Xenopus* egg extracts where DNA-coated beads can direct bipolar spindle assembly (Heald et al., 1997). Four molecular mechanisms have been proposed to drive chromosome-directed spindle assembly: the Ran-GTP pathway, the Chromosome Passenger Complex (CPC) pathway, the kinetochore pathway, and the Augmin pathway. These mechanisms have been summarized in (Bennabi et al., 2016).

The small GTPase Ran has been demonstrated to play a critical role in chromosome-directed spindle formation in addition to its role in nuclear transport (Carazo-Salas et al., 1999; Drutovic et al., 2020; Hetzer et al., 2002; Kalab et al., 1999). The Ran GEF, RCC1, which creates Ran-GTP by promoting exchange of GTP for GDP, is localized in interphase nuclei and on condensed chromatin from prometaphase through anaphase during mitosis in cultured human cells (Moore et al., 2002; Ohtsubo et al., 1989), cultured rodent cells (Li et al., 2003), and *Xenopus* sperm chromatin incubated in M-phase *Xenopus* egg extract (Bilbao-Cortés et al., 2002; Li et al., 2003). Ran-GTP is thus concentrated in close proximity to chromosomes at nuclear envelope breakdown where it can release inactive spindle assembly factor from binding with importins, thereby stimulating spindle assembly locally near chromosomes (Figure 1A). Cytoplasmic Ran-GAP converts Ran-GTP to Ran-GDP far from chromosomes, thus inhibiting spindle assembly in regions far from chromatin. In *Xenopus* egg extracts, Ran-GTP is necessary for spindle assembly around DNA-coated beads, constitutively active Ran-GTP stimulates spindle assembly in the absence of DNA-coated beads (Carazo-Salas et al., 1999) and RCC1 linked to beads is sufficient to drive bipolar spindle assembly (Halpin et al., 2011). The role of Ran-GTP in living oocytes has been less clear. An early study found that manipulating levels of Ran-GTP in mouse oocytes did not inhibit assembly of functional meiosis I spindles (Dumont et al., 2007). RCC1 is distributed throughout mitotic cytoplasm in *Drosophila* embryos (Frasch, 1991) and depletion of Ran-GTP did not abolish *Drosophila* meiotic spindle formation (Cesario and McKim, 2011). In contrast, Ran-GTP is required for meiotic spindle assembly in human oocytes (Holubcová et al., 2015). Later studies found that the requirement for Ran-GTP in mouse oocytes is redundant with pericentrin-dependent MTOCs (Baumann et al., 2017; So et al., 2022) whereas another study using a different method for inhibiting Ran-GTP found that it was required non-redundantly for meiotic spindle assembly in mouse oocytes (Drutovic et al., 2020). In *C. elegans*, depletion of Ran by RNAi prevented assembly of mitotic spindles (Askjaer et al., 2002) but did not prevent meiotic spindle assembly (Chuang et al., 2020).

**Fig 1.**
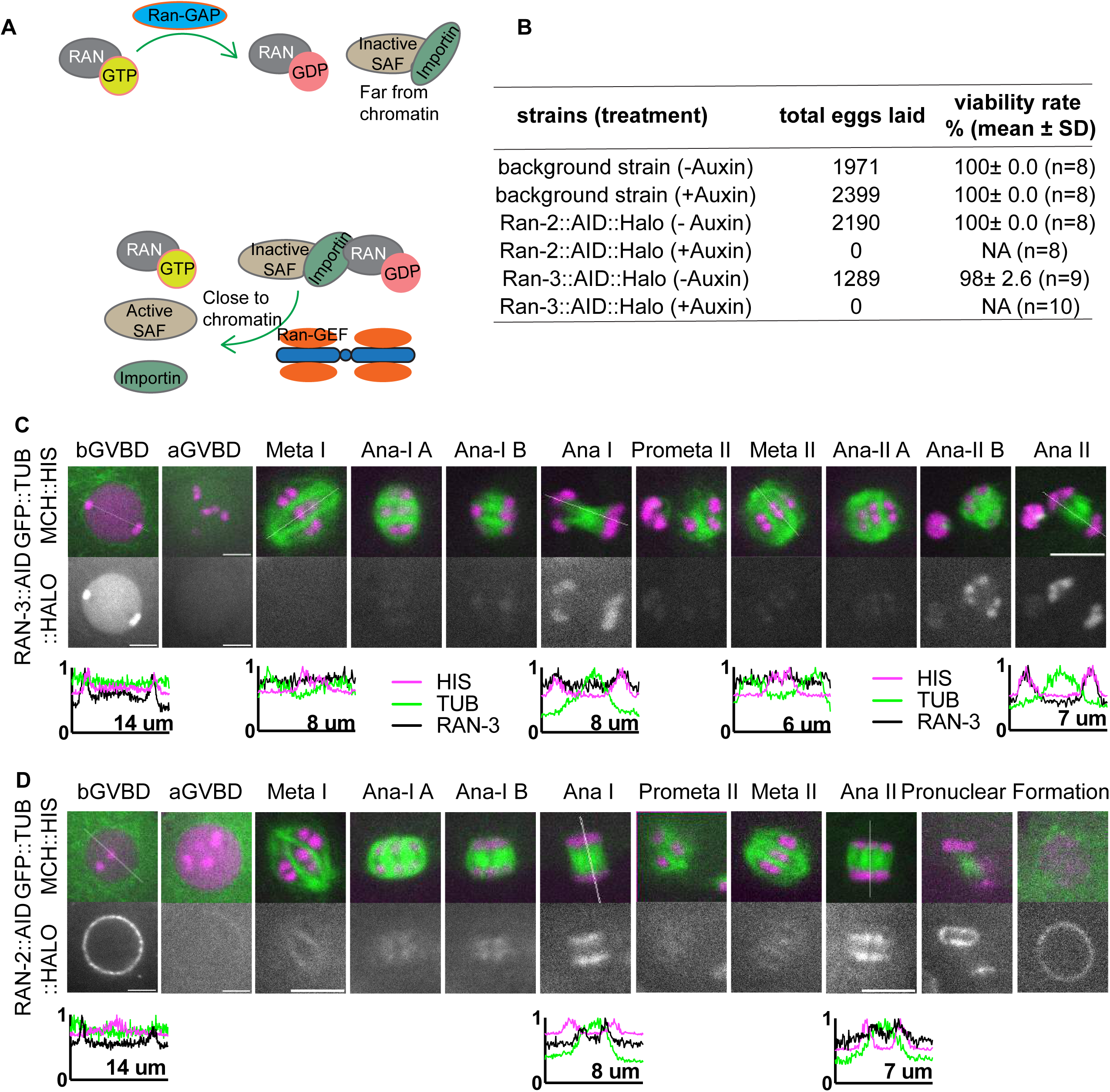
RAN-2 and RAN-3 are associated with chromosomes during meiotic anaphase. (A) Diagram of Ran-GDP, Ran-GTP cycle activating spindle assembly factors (SAFs) near chromatin. (B) Embryonic viability of strains after depleting RAN-2 or RAN-3 by auxin-induced degradation. (C) Time lapse images of meiotic embryo expressing endogenously tagged RAN-3::AID::HALO, TIR1::mRuby, GFP::TUB (tubulin) and mCh::HIS (mCherry::histone H2b). (D) Time lapse images of meiotic embryo expressing endogenously tagged RAN-2::AID::HALO, TIR1::mRuby, GFP::TUB (tubulin) and mCh::HIS (mCherry::histone H2b). Graphs representing normalized fluorescence intensity along a 1-pixel-wide line scan (indicated by dashed line on the respective images) before GVBD, Ana I and Ana II are plotted on the bottom. Scale bars, 5μm.

The CPC is composed of Aurora B/C kinase, the inner centromeric protein (INCENP), Survivin, and Borealin, which target it to chromatin. The CPC is required for bipolar meiotic spindle assembly in mouse (Nguyen et al., 2018) and *C. elegans* (Divekar et al., 2021) although a cloud of disorganized microtubules is still nucleated near chromosomes. The Augmin complex is activated directly (Kraus et al., 2023) and indirectly (Petry et al., 2013) by Ran-GTP and recruits γ-tubulin to nucleate new microtubules on the sides of preexisting microtubules (Colombié et al., 2013; Goshima et al., 2008; Lawo et al., 2009; Petry et al., 2011; Sánchez-Huertas and Lüders, 2015; Uehara et al., 2009). However, no augmin homologs have been reported in the *C. elegans* genome.

A less-studied centrosome-independent pathway that may promote spindle assembly in the vicinity of chromosomes is the concentration of α/β-tubulin dimers in the nuclear volume at Nuclear Envelope Breakdown (NEBD). Unpolymerized tubulin, monitored in cells treated with microtubule-depolymerizing drugs, is excluded from nuclei during interphase. During NEBD, rather than equilibrating to equal concentrations in the cytoplasm and nucleus, tubulin dimers have been reported to concentrate in the nuclear volume in *Drosophila* mitotic embryos (∼ 1.6-fold, Yao et al., 2012), *C. elegans* mitotic embryos (∼ 2-fold, Hayashi et al., 2012), *Drosophila* S2 cells (∼ 1.5-fold, Schweizer et al., 2015) and *Drosophila* neuroblasts (Métivier et al., 2021). The concentration of tubulin dimer independently of microtubule polymerization has been proposed to be related to binding to a spindle matrix, however, non-proteinaceous molecules like dextran (a polysaccharide) can also concentrate in the nuclear region (Yao et al., 2012), raising the question whether binding to a spindle matrix is necessary or not. Depletion of Ran by RNAi affects this concentration and causes spindle defects in *C. elegans* mitotic embryos (Hayashi et al., 2012) and *Drosophila* neuroblasts (Métivier et al., 2021). It is unclear whether the concentration of tubulin may be a component of the Ran pathway, or the defects observed might be the indirect result of altering the kinetics of nuclear import/export long before mitosis.

*C. elegans* female meiosis is unique in a few different ways. First, no distinct MTOCs containing PCM proteins have been observed in mature oocytes or meiotic embryos (McNally et al., 2006; Wolff et al., 2016). Second, RNAi knockdown of *C. elegans* Ran, RAN-1, decreases spindle microtubule levels but does not block meiotic spindle formation (Chuang et al., 2020). Third, *C. elegans* Katanin, composed of MEI-1 and MEI-2, is concentrated on chromosomes and is essential for formation of meiotic spindle poles (McNally et al., 2014; Srayko et al., 2000). Fourth, depletion of *C. elegans* γ-tubulin, TBG-1, by RNAi also leads to spindle microtubule loss but does not prevent meiotic division, although oocytes depleted of γ-tubulin and katanin by RNAi assemble extremely reduced levels of microtubules around chromosomes (McNally et al., 2006). Lastly, most SAFs remain cytoplasmic prior to GVBD (McNally et al., 2022). This suggests there might be novel mechanisms underlying the microtubule nucleation and spindle assembly without centrosomes in *C. elegans* oocytes. The mechanisms observed in *C. elegans* may be conserved in mammals and our studies may reveals factors that could be potential therapeutic targets to improve the efficacy of IVF.

## Results

### RAN-2^Ran-GAP^ and RAN-3^Ran-GEF^ are enriched on anaphase chromosomes

A previous study depleting Ran in *C. elegans* by *ran-1(RNAi)* showed female meiotic spindles still formed although with a reduced density of microtubules and showed relatively normal anaphase progression (Chuang et al., 2020). As RAN-1 may not have been fully depleted by RNAi, we created conditional knockdown worm strains of Ran-GEF and Ran-GAP by adding Auxin Induced Degron (AID) (Zhang et al., 2015) and HALO tag sequences to the endogenous *ran-2*^Ran-GAP^ and *ran-3*^Ran-GEF^ loci. RAN-2::AID::HALO and RAN-3::AID::HALO worms laid no eggs on auxin plates (Figure 1B). Previous depletion of RAN-3 or RAN-2 by RNAi caused >90% embryonic lethality but did not affect brood size (Askjaer et al., 2002). This suggests that depletion of RAN-2 or RAN-3 through the AID system results in more complete depletion than RNAi.

It has been proposed that chromosome associated RCC1(Ran-GEF) generates Ran-GTP near chromatin while cytoplasmic Ran-GAP generates Ran-GDP far from chromosomes. Ran-GTP proximal to chromosomes releases inactive SAFs from binding with importins (Figure 1A). Consistent with these ideas, we observed RAN-3::AID::HALO in the nucleoplasm and chromosomes before GVBD (Figure 1C). After GVBD, RAN-3 diffused from the nucleus and was not detected on chromosomes during spindle assembly in 12/12 embryos. RAN-3 only faintly associated with chromosomes at metaphase I and II. In contrast, RAN-3 strongly localized to chromosomes at anaphase I and anaphase II (Figure 1C). RAN-2::AID::HALO was strongly associated with the nuclear envelope before GVBD in 11/11 oocytes. Later, RAN-2 faintly labeled metaphase I and metaphase II spindles. At anaphase, RAN-2 localized to the spindle midzone, and its intensity increased as anaphase progressed. It is strongly associated with the inner side of separating chromosomes (Figure 1D). These results suggest that RAN-3 and RAN-2 might function primarily at anaphase.

### Ran-GEF is required for extrusion of the first polar body but not metaphase I spindle formation

Ran regulators are also involved in nuclear transport, defects in which usually lead to small and leaky nuclei. By treating the worms with auxin for a brief period, we sought to only evaluate the function of RAN-3 and RAN-2 on meiotic spindle formation without disrupting meiotic prophase and nuclear transport. When treated with auxin for 4 hours, expression of RAN-3::AID::HALO in the nucleoplasm and chromosomes in diakinesis oocytes was significantly reduced (Figure S1A-B). At 6-hour auxin treatment, the expression level was comparable to control oocytes with no HALO expression. Moreover, -1 oocytes treated with auxin for 4 hours or 6 hours were smaller than controls, whereas 36-hour auxin resulted in much smaller nuclei (Figure S2A, B). Qualitatively, GFP:: tubulin partially leaked from the cytoplasm into nuclei after 4 hours of auxin but GFP::tubulin leaking into the nuclei was much more severe at 36 hours of auxin (Figure S2C). These results indicated that RAN-3 depletion after 4 hours of auxin results in pre-NEBD defects and defects are more severe after 36 hours of auxin. Expression of RAN-2::AID::HALO in the nucleoplasm and nuclear envelope in diakinesis oocytes was reduced to control levels after 4 hours of auxin (Figure S1C, D). The sizes of oocyte nuclei were normal in RAN-2::AID::HALO worms after 4 or 6 hours of auxin, and their nuclei were not leaky compared to controls (Figure S2A-C). RAN-2::AID::HALO oocytes treated with auxin for 36 hours were not quantified as these oocytes were severely disorganized. We therefore analyzed meiotic spindle assembly by time-lapse imaging after 4 to 6 hours of auxin because these timepoints showed strong depletion but no detectable effect on nuclear function for RAN-2 and only moderate early effects for RAN-3.

Bipolar metaphase I spindles were observed in control worms (n= 15 no degron plus auxin; n= 10 *ran-3::AID* no auxin; n=5 ran-2::AID no auxin) (Fig. 2A; Video 1). Consistent with the dispersal of RAN-3 and RAN-2 at nuclear envelope breakdown (Figure 1C, D), 4 to 6-hour auxin treatment of RAN-2::AID::HALO (n = 33) or RAN-3::AID::HALO (n = 26) worms (Figure 2B, C) also resulted in bipolar metaphase I spindles. The mean metaphase I spindle GFP::tubulin pixel intensities were significantly decreased after RAN-3 depletion relative to both no degron controls or no auxin controls (Fig. 2E, F). The length of metaphase I spindles was not significantly different than controls after depletion of RAN-2 or RAN-3 (Fig. S1E) although a slight increase in metaphase II spindle length was observed after RAN-2 depletion (Fig. S1F). The velocity of anaphase I (reported as the rate of increase in distance between separating chromosome masses) was significantly faster than no degron or no auxin controls after depletion of RAN-3 (Fig. 2G). However, the velocity of anaphase II was not significantly different than controls (Fig. S1G).

**Fig 2.**
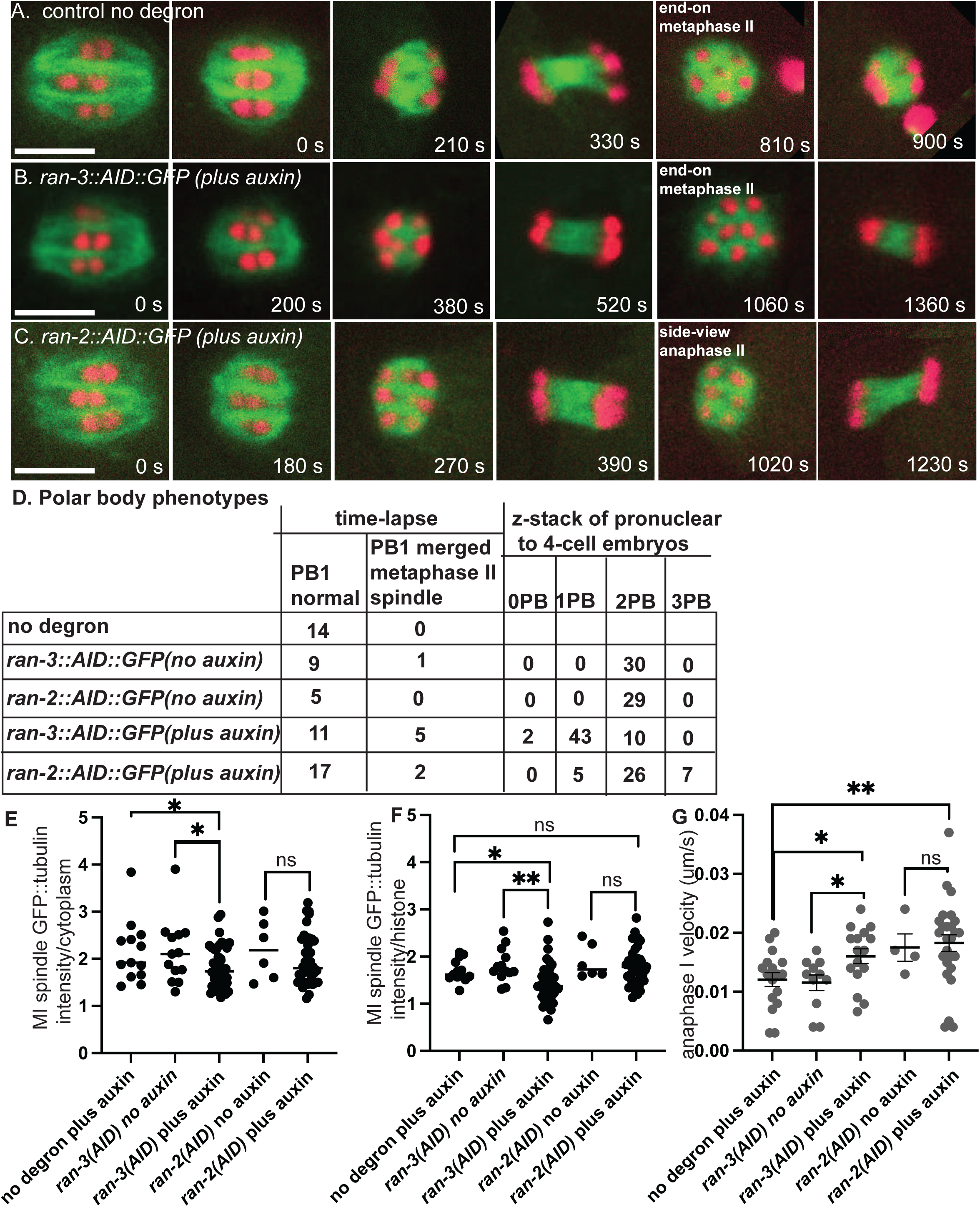
Ran-GEF is required for extrusion of the first polar body but not for meiotic spindle formation. (A-C) Representative images from single focal plane, in utero time-lapse imaging from metaphase I through anaphase II of worms treated with auxin for 4 - 6 hrs. Metaphase I spindle morphology was not affected by depletion of RAN-2 or RAN-3. (A) At 810s, 6 chromosomes are visible in an end-on view of a metaphase II control spindle. (B) At 1060 s, 8 chromosomes are visible in an end-on view of a RAN-3-depleted metaphase II spindle, indicating failure of the first polar body. (C) At 1020 s, 3 chromosomes are visible in a side-on view of an early anaphase II RAN-2-depleted spindle. (D) Number of embryos with different polar body phenotypes interpreted from either single plane time-lapse imaging or z-stacks of 1-4 cell post-meiotic embryos. (E) Mean GFP::tubulin pixel intensity of the entire metaphase I spindle divided by the mean GFP::tubulin intensity adjacent to the spindle. (F) Mean GFP::tubulin pixel intensity of the entire metaphase I spindle divided by the mean mCherry::histone H2b intensity of the brightest half bivalent. (G). Anaphase I velocities measured as the increase in distance between separating chromosome masses divided by time. Bar = 5 um. * p< .05, ** p < .01.

In 5/16 time-lapse sequences of RAN-3-depleted worms, chromosomes that moved toward the cortex at anaphase I merged into the metaphase II spindle. When viewed end-on, control metaphase II spindles have 6 univalents arranged in a pentagonal array (Fig. 2A, 810 s). In a side view of this pentagonal array during metaphase or early anaphase, 3 chromosome pairs are in focus (Fig. 2C, 1020 s). Merging of polar body-bound anaphase I chromosomes into the metaphase II spindle was inferred from more than 6 chromosomes in an end-on view (Fig. 2B, 1060 s) or from direct observation when merging occurred in a favorable focal plane (Video 2). However, this frequency of polar body failure was not significantly different than the 1/10 failures observed in no auxin controls of the same strain (Fig. 2D; p=.35 Fisher’s exact test). Thus failure to extrude the first polar body might be an artifact of time-lapse imaging or the genetic background. Conversely, the actual polar body failure rate might be much higher if the chromosomes that segregated toward the cortex at anaphase I do not move inward until later in development. To address both issues, we collected z-stacks of post-meiotic pronuclear to 4-cell stage embryos dissected from worms at 5-7 hrs after placing worms on auxin. These embryos underwent meiosis 4-6 hrs after auxin treatment and the worms were not mounted for time-lapse imaging when these embryos underwent meiosis. 43/55 RAN-3 depleted embryos had a single polar body, whereas 29/29 no auxin controls of the same strain had 2 polar bodies (Fig. 2D). This difference was significant (p=.0001 Fisher’s exact test). Because the chondroitin layer of the eggshell is secreted during anaphase I (Olson et al., 2012), the first polar body is embedded in the eggshell and remains stationary at the anterior tip of the oval embryo, while the second polar body is inside the eggshell and moves to an internal location over time. Consistent with this, 18/19 no auxin control 1-cell embryos had 2 polar bodies at the anterior tip whereas 19/20 no auxin control 4-cell embryos had one polar body at the tip and one polar body internal. The single polar body of 14/15 RAN-3 depleted 1-cell embryos was at the anterior tip whereas the single polar body of 15/15 RAN-3 depleted 4-cell embryos was internal. The 4-cell result was significantly different than a random distribution of 7/7 (p= .002 Fisher’s exact test) and suggests that RAN-3 depletion causes resorption of chromosomes destined for the first polar body. Polar body extrusion defects after RAN-2 depletion compared with the same strain with no auxin were less definitive (p = .052 Fisher’s exact test) and chromatin was extremely condensed in RAN-2 depleted post meiotic embryos, making it difficult to distinguish polar bodies from micronuclei at the cortex. Overall, these results suggest that the Ran-GTP pathway is important for limiting anaphase I velocity and polar body formation. Notably, RAN-3 is concentrated on chromosomes during anaphase and polar body extrusion (Fig. 1). RAN-3 may have a more limited role in promoting the density of spindle microtubules at metaphase I when its localization on chromosomes is not discernible (Fig. 1).

### Free tubulin is concentrated in the nuclear volume at GVBD

At the onset of mitosis in *C. elegans* and *Drosophila*, soluble tubulin concentrates in the nuclear/spindle volume relative to the surrounding cytoplasm (Hayashi et al., 2012; Métivier et al., 2021; Schweizer et al., 2015; Yao et al., 2012). Since microtubule polymerization is concentration dependent, this concentration might facilitate spindle formation in the vicinity of chromosomes in meiotic oocytes. In DMSO-treated control -1 oocytes, mNG::TBB-2 (β-tubulin) labelled microtubules in the cytoplasm and was excluded from the nucleus (Figure 3A, 3B). Upon fenestration of the nuclear envelope (GVBD: germinal vesical breakdown), indicated by leakage of non-chromosomal mCherry::histone out of the nucleus, mNG::TBB-2 fluorescence increased within the nuclear volume in 10/10 time-lapse sequences (Figure 3B) as previously described (McNally et al., 2006; Mullen and Wignall, 2017). Tubulin fluorescence then transformed into a “microtubule cage” (Mullen and Wignall, 2017) and eventually a bipolar spindle in 10 out of 10 time-lapse sequences. In oocytes treated with nocodazole to depolymerize microtubules, tubulin was diffuse in the cytoplasm prior to GVBD (Figure 3C), and still concentrated in the nuclear volume during and after GVBD (Figure 3C; Figure 3D, 3E). Simple diffusion from the cytoplasm into the nuclear volume should result in equal fluorescence intensities of mNG::TBB-2 in the nucleus and cytoplasm but fluorescence instead increased in the nuclear volume to 1.2-fold greater than the cytoplasm (Figure 3E). After ovulation, chromosomes in nocodazole-treated zygotes were dispersed and loosely wrapped by sparse short microtubules. No spindle formation or chromosome separation was observed before pronucleus formation in 12 out of 12 time-lapse sequences.

**Fig 3.**
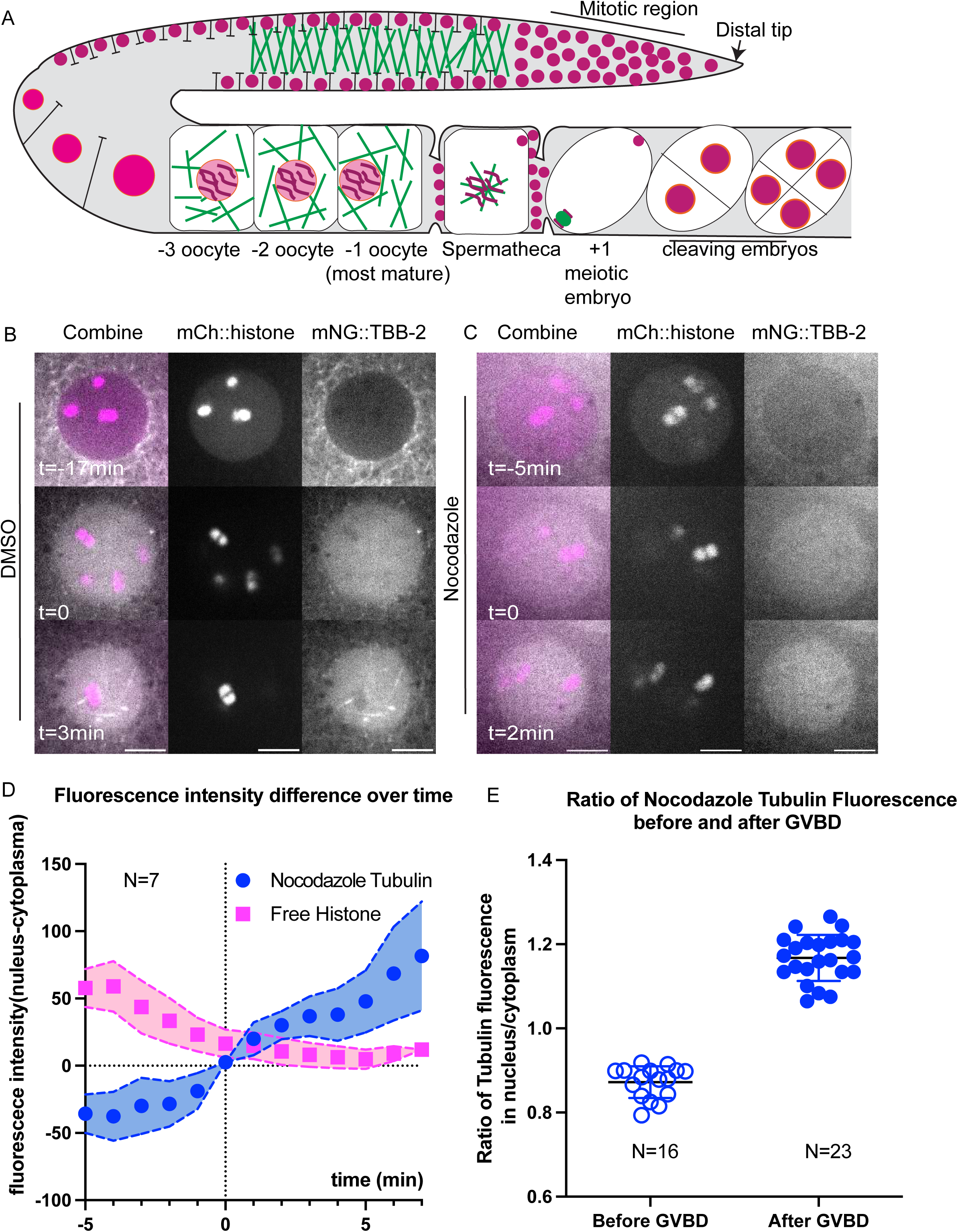
Unpolymerized tubulin concentrates in the nuclear volume at germinal vesicle breakdown (GVBD). (A) Diagram of chromosome and microtubule organization in the *C. elegans* gonad. DNA (magenta); Microtubules (Green); Plasma Membrane (Black); Nuclear Envelope (Orange); -1 oocyte: most mature prophase-arrested oocyte; +1 embryo: fertilized embryo undergoing meiotic divisions. (B, C) Representative time-lapse images of the Germinal Vesicle in -1 oocyte expressing mNG::TBB-2 (mNeonGreen::tubulin: greyscale) and mCh::histone (mCherry::histone H2b: magenta). Tubulin concentrated in the nuclear volume at GVBD in worms treated with (B) DMSO or (C) Nocodazole. Non-chromosomal histone in the “nucleus” diffuses out at the onset of GVBD. bGVBD, GVBD, aGVBD: before, at and after GVBD. Scale bars, 5 μm. (D) Plots of fluorescence intensity difference in nucleus and cytoplasm over time after nocodazole treatment. Tubulin (blue); Histone (Magenta). Y axis: fluorescence intensity in nucleus - fluorescence intensity in cytoplasm. N: number of time lapse sequences analyzed. Mean shown in solid squares [His] or solid circles [Tub]. SEM shown in colored regions. (E) Ratio of mean fluorescence intensity of nocodazole-tubulin in nucleus over cytoplasm before GVBD and after GVBD. Tubulin were excluded from nucleus before GVBD (ratio < 1) and concentrated in nucleus after GVBD (ratio > 1). N: number of nuclei analyzed.

We noticed that before GVBD, fluorescence of mNG::TBB-2 was detectable in the nucleus, which contradicts with the general assumption that the nucleus is void of tubulin (Figure 3B top panel, S2B, average of the ratio of tubulin fluorescence in the nucleus to the cytoplasm: 0.69). To determine whether there is tubulin in the nucleus or if this is due to pinhole crosstalk from the spinning disk confocal, we captured images on a Zeiss laser scanning confocal microscope which has a single pinhole and therefore removes out of focus light more efficiently. The ratio of tubulin fluorescence in the nucleus to the cytoplasm before GVBD was significantly reduced to 0.33 (Figure S3A-B) in images from the Zeiss LSM compared with 0.69 from the spinning disk confocal. The difference in nuclear vs cytoplasmic fluorescence was also greater on the Zeiss LSM for other probes (Fig. S3C-D). This suggests that the actual concentration difference between nucleus and cytoplasm are likely greater than the ratios reported from spinning disk confocal images.

Similar results in previous studies of mitosis led to the interpretation that alpha/beta tubulin dimers concentrate in the nuclear volume of unperturbed cells during spindle assembly. However, it is possible that nocodazole does not completely block microtubule polymerization and the fluorescence accumulating in the nuclear volume of nocodazole-treated oocytes represents accumulation of short microtubules. It is also possible that this phenomenon is induced by nocodazole and does not occur in unperturbed cells.

### Accumulation of tetrameric GFP and un-polymerizable tubulin in the “nuclear volume” at GVBD

Previous investigators suggested that tubulin dimers concentrate either by binding to something in the nuclear volume (Hayashi et al., 2012; Métivier et al., 2021) or by being excluded by cytoplasmic organelles that are kept out of the spindle volume by the ER envelope that still envelopes the spindle after nuclear envelope breakdown (Schweizer et al., 2015; Figure 4A). We analyzed the behavior of two fluorescent probes designed to address three major issues: 1. Incomplete depolymerization by nocodazole; 2. Concentration of alpha/beta dimers in the absence of nocodazole; 3. Specific binding of tubulin to a nuclear binding site vs volume exclusion by cytoplasmic organelles. GFP::GCN4-pLI is a tetramerized GFP designed to have a native molecular weight (128 kD) similar to an alpha/beta tubulin dimer (101 kD without tag, 128 kD with mNeonGreen tag, see Fig. 7A), but which should not bind to any tubulin-specific binding sites in the nuclear volume because GCN4-pLI is a well characterized synthetic 4 helix bundle (Mittl et al., 2000). GFP::TBA-2(T349E) is an alpha tubulin mutant that can dimerize with beta tubulin, cannot polymerize (Johnson et al., 2011), and should bind any specific tubulin binding sites in the nuclear volume. Both GFP::GCN4-pLI (Figure 4B; Video 4) and GFP::TBA-2(T349E) (Figure 4C; Video 5) concentrated in the nuclear volume at GVBD in the absence of nocodazole. These results 1. Suggested that the concentration of mNG::TBB-2 in nocodazole was not due to incomplete depolymerization; 2. That tubulin dimers concentrate in the nuclear volume during unperturbed spindle assembly; and 3. That binding to a tubulin-specific binding site in the nuclear volume is not required for concentration.

**Fig 4.**
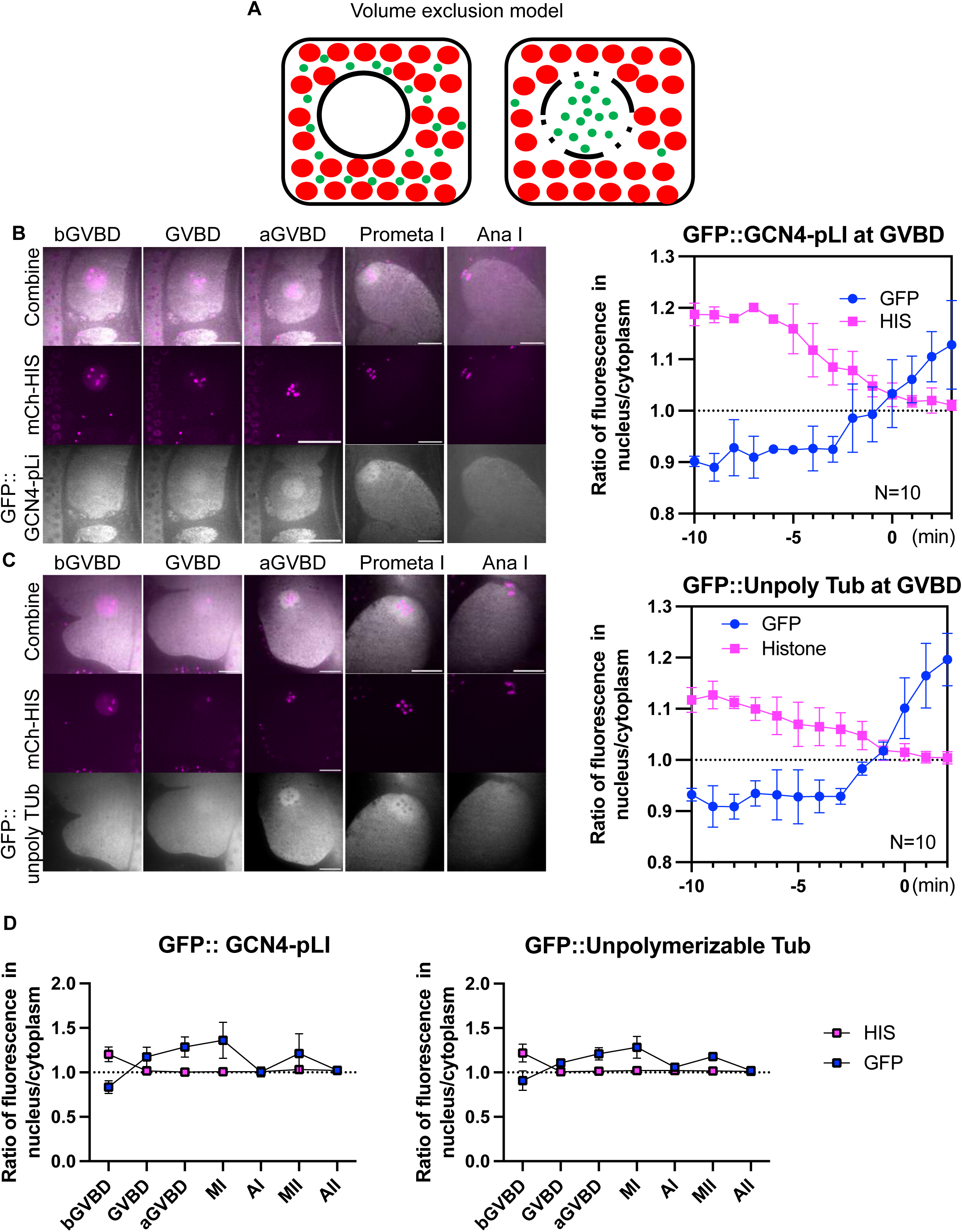
Tubulin-sized molecules concentrated in the nuclear volume at GVBD. (A) Volume exclusion model of free tubulin rushing into nucleus (black circled region) where more space is available to tubulin-sized molecules (green dots) at germinal vesicle breakdown (GVBD) due to volume occupied by mitochondria and yolk granules (red dots). (B) Representative time lapse images of meiotic embryo expressing tetrameric GFP::GCN4-pLI (greyscale) and mCh::HIS (mCherry::histone H2b: magenta). The ratio of fluorescence intensity of GFP::GCN4-pLI or non-chromosome histone in nucleus to cytoplasm during GVBD over time is shown in the graph on the right. N: number of time lapse sequences analyzed. Means are shown in solid square [His] or solid circle [Tub]. Bars indicate SEM. Scale Bars, 10um. (C) Representative time lapse images of a meiotic embryo expressing un-polymerizable GFP::TBA-2(T349E) (greyscale) and mCh::histone (magenta). The ratio of fluorescence intensity of GFP or non-chromosome histone in nucleus to cytoplasm during GVBD over time is shown in the graph on the right. Scale Bars, 10um. (D) Plots of fluorescence intensity ratio in the nucleus or spindle to cytoplasm before germinal vesicle breakdown (bGVBD), GVBD, or after GVBD (aGVBD), metaphase I (MI), anaphase I (AI), metaphase II (MII) and anaphase II (AII) show concentration at MI and MI, and dispersion at AI and AII.

Because GFP::GCN4-pLI and GFP::TBA-2(T349E) could be tracked in the absence of nocodazole, we could examine their behavior during normal meiotic divisions. Chromosome segregation was normal in GFP::TBA-2(T349E) oocytes and GFP::GCN4-pLI oocytes (12/12 and 10/10 filmed respectively) suggesting that expression of GFP::TBA-2(T349E) or GFP::GCN4-pLI did not severely disturb normal spindle function. Interestingly, their concentration at GVBD lasted through Metaphase I but diffused to a 1:1 spindle: cytoplasm ratio at Anaphase I, followed by re-accumulation at metaphase II and dispersion at Anaphase II (Figure 4B, 4C, 4D; Video 3, 5). This raises the question of what barrier between spindle and cytoplasmic volume might change between anaphase I and metaphase II.

### The ER delimits the accumulation of free tubulin during meiosis and early mitosis

Re-accumulation of tubulin within the metaphase II spindle volume could be due to nuclear import if the nuclear envelope transiently reformed or the re-accumulation could be due to cell-cycle changes in the structure of the ER that surrounds the spindle (Fig. 5A; Poteryaev et al., 2005). To test whether a functional nuclear envelope reforms transiently between anaphase I and metaphase II, we tracked GFP::NPP-6, a nuclear pore protein of the Y complex (Galy et al., 2003) (Figure 5B) and GFP::LMN-1, the *C. elegans* nuclear Lamin (Galy et al., 2003) (Figure 5C) by time-lapse imaging. Both proteins disappeared from the nuclear envelope at GVBD and did not re-locate to the nuclear membrane until pronuclear formation (Figure 5B, C). The absence of NPP-6 from a nuclear envelope during meiosis has been previously reported (Penfield et al., 2020). These results make it unlikely that tubulin concentrates in the spindle volume at metaphase II due to nuclear import. In contrast, the ER is contiguous with the outer nuclear membrane before GVBD (Figure 5A, 5D). After GVBD, the ER wraps around meiotic spindles with clustering at spindle poles during metaphase I and metaphase II (Figure 5D). In contrast, yolk granules (Figure 5D), maternal mitochondria (Figure 5E), and lipid droplets (Fig. 5F) are excluded from the nuclear and spindle region. At the light microscope level (Figure 5D), the ER appears reticular during metaphase I and metaphase II and disperses during anaphase I and anaphase II as described previously (Poteryaev et al., 2005). To determine the ultrastructural changes in ER morphology, we manually segmented the ER in previously published electron tomograms (Lantzsch et al., 2021). During metaphase I and metaphase II, the sides of the spindle are enclosed by overlapping sheets of ER (Video 6, Figure 6A; Video 7) and the accumulation of ER at the spindle poles is a complex mixture of sheets and tubules (Figure 6B; Video 8). During anaphase I and anaphase II, the ER near the spindle consists entirely of tubules (Figure 6C, 6D; Video 9). By time-lapse imaging of worms expressing an ER marker as well as GFP::GCN4-pLI, we found that the dramatic morphological change of ER coincides with the concentration and dispersion of GFP::GCN4-pLI during meiosis (Video 3) and mitosis (Video 10). Thus tubulin-sized proteins accumulate in the spindle volume when it is encased by sheet-like ER and disperse when the ER is tubular. These results favor a model in which a semi-permeable spindle envelope composed of sheet-like ER excludes a “crowding agent” from the spindle volume, so that tubulin-sized proteins concentrate in the spindle volume by volume exclusion from the surrounding cytoplasm (Figure 4A).

**Fig 5.**
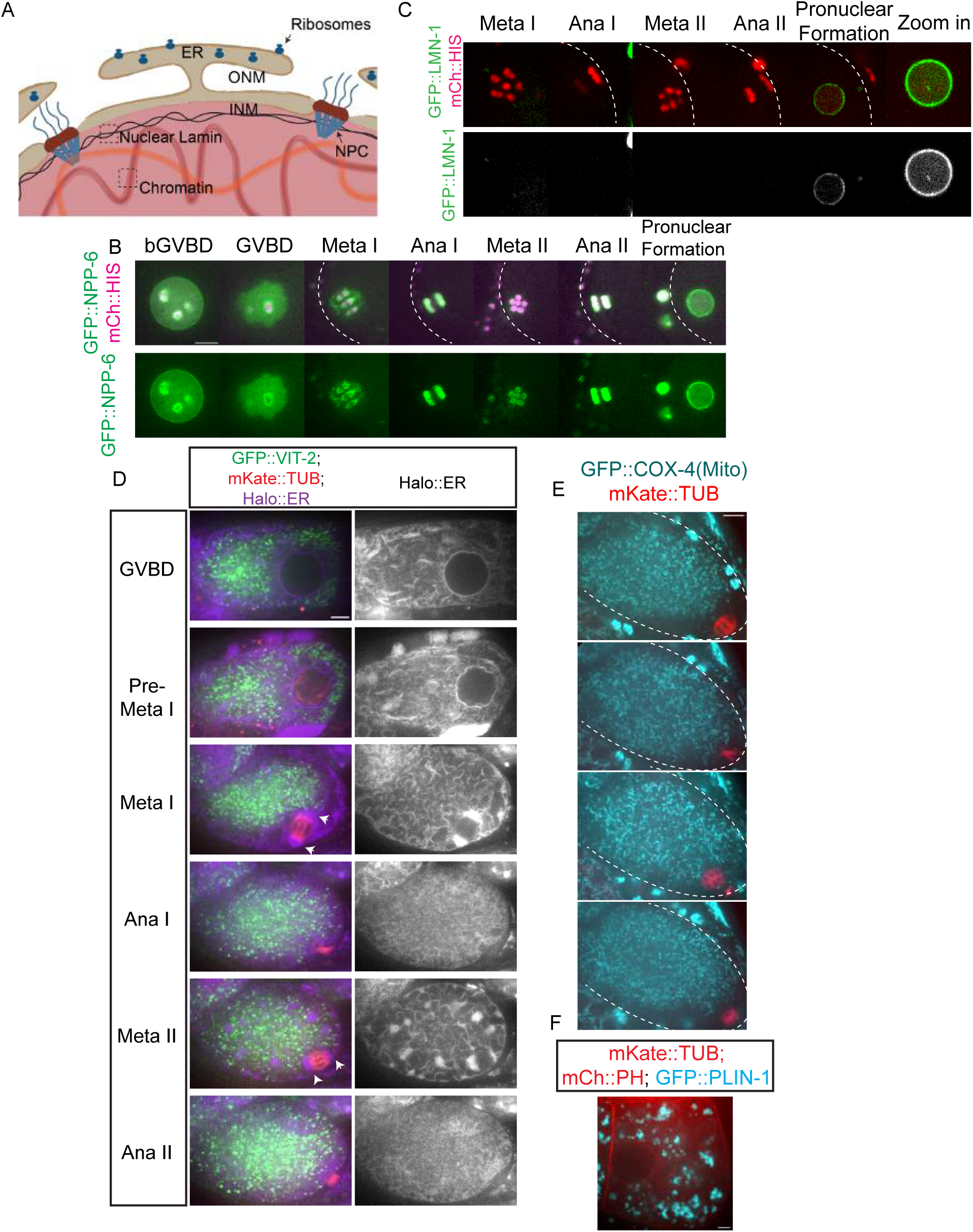
The ER envelope delimits the concentration of free tubulin during meiosis. (A) Diagram of an intact nucleus before GVBD. ONM: Outer Nuclear Membrane. INM: Inner Nuclear Membrane. NPC: Nuclear Pore Complex. ER: Endoplasmic Reticulum. (B) Time-lapse sequences of representative meiotic embryo expressing GFP::NPP-6 (green); mCh::histone (red) (C) or GFP::LMN-1 (green); mCh::histone (magenta) (D) or GFP::VIT-2 (yolk granule cargo: green); mKate::TUB (red); HALO::ER (magenta) (E) or GFP::COX-4 (mitochondrial protein: cyan); mKate::TUB (red) (F) or GFP::PLIN-1 (lipid droplet protein: cyan); mCh::PH (red); mKate::TUB (red). Scale Bars = 5um. The cell cortex was drawn in white dashed line in B, C and E.

**Fig 6.**
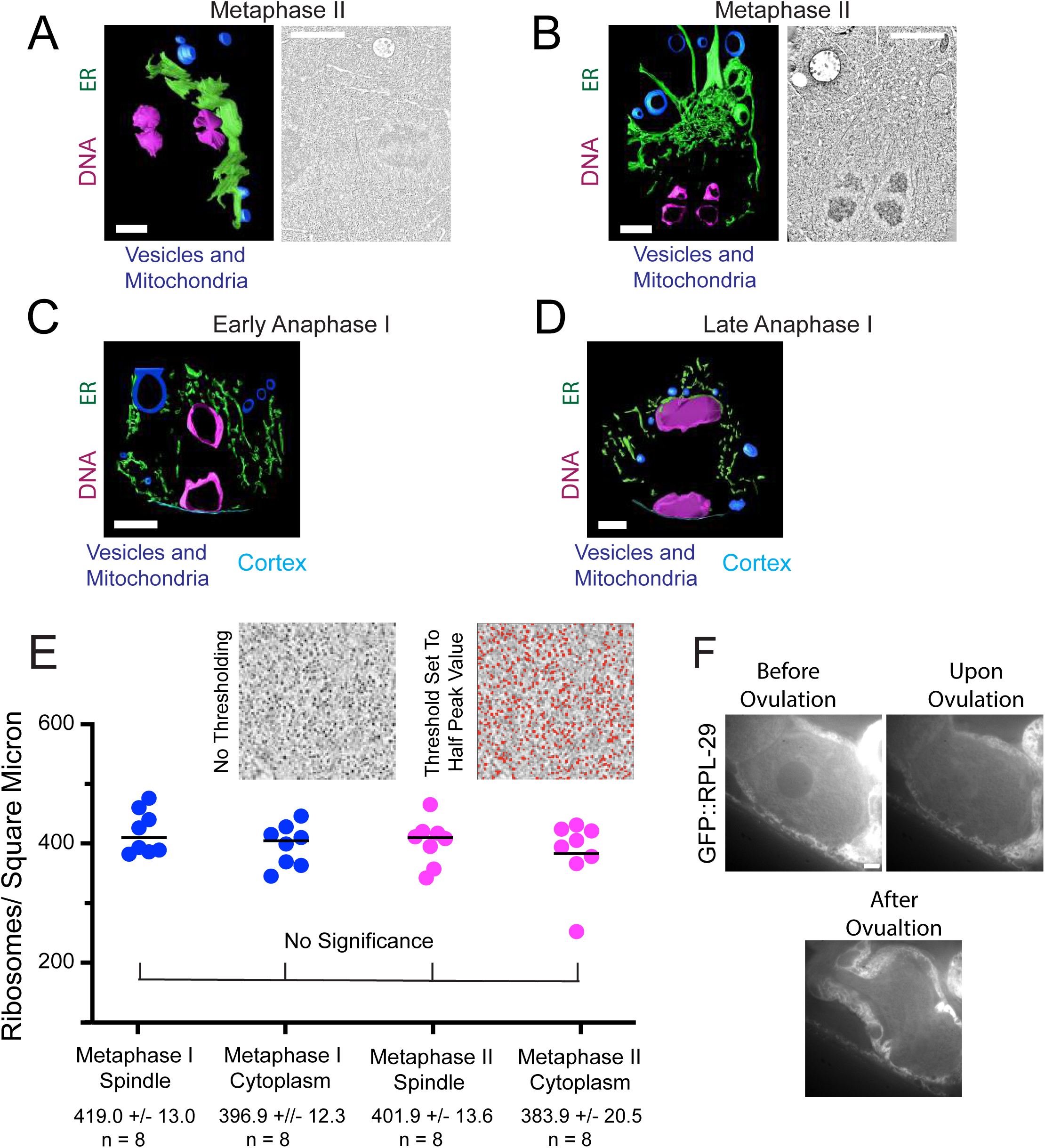
ER sheets envelop metaphase meiotic spindles. (A) Left. Model of ER sheets on the partial exterior of a metaphase II meiotic spindle derived from an electron tomogram and spanning 1.2 um in the z. Right. Single plane image from the tomogram. The 2 lobes of each univalent chromosome are oriented down the pole to pole axis of the spindle. (B) Left. Model of ER at one pole of a metaphase II meiotic spindle derived from an electron tomogram and spanning 0.6 um in the z. Right. Single plane image from the tomogram. The 2 lobes of each univalent chromosome are oriented down the pole to pole axis of the spindle. (C) Model of ER in an outer 0.6 um z-section of a MI early anaphase spindle. No large ER sheets were observed. (D) Model of ER in a 0.6 um z-section of a late anaphase I spindle. (A-D) ER in green, chromosomes in magenta, cytoplasmic organelles in blue. (A-D) Scale bar, 1 um. (E) Ribosomes were counted in representative squares in representative sections of spindle and cytoplasm in metaphase I and metaphase II electron tomograms. Ribosomes were counted after setting the threshold value to half of the peak value. Average ribosome areas in metaphase I spindles and cytoplasm, and metaphase II spindles and cytoplasm were: 30.9 +/- 2.0nm, 28.0 +/- 1.2nm, 26.3 +/- 1.0nm, and 26.8 +/- 1.7nm respectively. (F) Time-lapse images of meiotic embryo expressing GFP::RPL-29 (grayscale) during GVBD. Scale Bar, 5um.

The “crowding agent” might consist of membranous organelles like yolk granules (Figure 5D), or mitochondria (Figure 5E) as suggested by Schweizer et al., 2015 or might consist of ribosomes as suggested by Delarue et al., 2018. To test whether ribosomes are a viable candidate for excluding volume from the cytoplasm, we determined the density of ribosomes inside of the spindle and outside the ER envelope from previously published electron tomograms (Lantzsch et al., 2021) and found no significant difference in ribosome density inside vs outside the spindle (Figure 6E). As a complementary approach, we monitored a ribosomal protein, GFP::RPL-29, by time-lapse imaging before and after GVBD (Figure 6F). GFP::RPL-29 was excluded from the nucleus before GVBD, then rapidly rushed into the nuclear volume at GVBD in 9/9 time-lapse sequences, indicating that ribosomes could not be the cytoplasmic crowding agent causing concentration of tubulin-sized proteins that stay concentrated through metaphase.

To test the idea that yolk granules, mitochondria, and other organelles are the crowding agent that excludes soluble tubulin from the cytoplasm, we segmented all the membranous organelles visible in previously described electron micrographs (Bembenek et al., 2007; Howe et al., 2001; Fig. S4A-D). In one thin section of a -1 oocyte, membranous organelles occupied 25% of total non-nuclear area. In two different sections covering an entire metaphase II embryo, organelles occupied 21% and 22% of total area. Membranous organelles occupied 28% and 31% of the non-spindle area in these sections. The reduced cytoplasmic volume outside the nucleus or metaphase spindle would thus be between 79% and 69%. The expected apparent concentration of molecules in the nuclear/spindle volume would thus be 1/.79 – 1/.69 or 1.3 – 1.4-fold. This is reasonably close to the observed 1.2-fold enrichment given the limitations of both measurements.

Notably, GFP::RPL-29 did not concentrate in the nuclear volume at GVBD but instead equilibrated to equal fluorescence intensity inside and outside the spindle envelope. This result indicated that there is specificity to the types of molecules that concentrate in the nuclear/spindle volume.

### The concentration of molecules during GVBD is size dependent

In immature starfish oocytes, fluorescent dextrans of 25 kDa or larger are excluded from the nucleus (Lénárt et al., 2003), similar to our results with mNG::TBB-2, GFP::GCN4pLI and GFP::TBA-2(T349E), presumably because they are too large to diffuse freely through NPCs, and/or because of nuclear export of tubulin (Schwarzerova et al., 2019). In contrast, 10 kDa fluorescent dextrans accumulated in the nucleus of immature starfish oocytes at a concentration twice that of the cytoplasm (Lénárt et al., 2003). It was suggested that this is because the small dextrans diffuse freely through NPCs and because yolk granules occupy 50% of the cytoplasmic volume thus driving apparent concentration of small dextrans in the nucleus (Lénárt et al., 2003). We found that a 36 kDa monomeric GFP concentrated in *C. elegans* oocyte nuclei before GVBD (Figure 7A, B) to a concentration twice that of the cytoplasm (Figure 7D), like 10 kDa dextrans in starfish oocytes. To rule out the possibility that this might be mediated by a cryptic NLS on GFP, we expressed a 34 kDa monomeric HALO tag in the *C. elegans* germline, which also concentrated to a 2-fold higher concentration in the nucleus relative to the cytoplasm in diakinesis oocytes prior to GVBD (Figure 7C-E).

**Fig 7.**
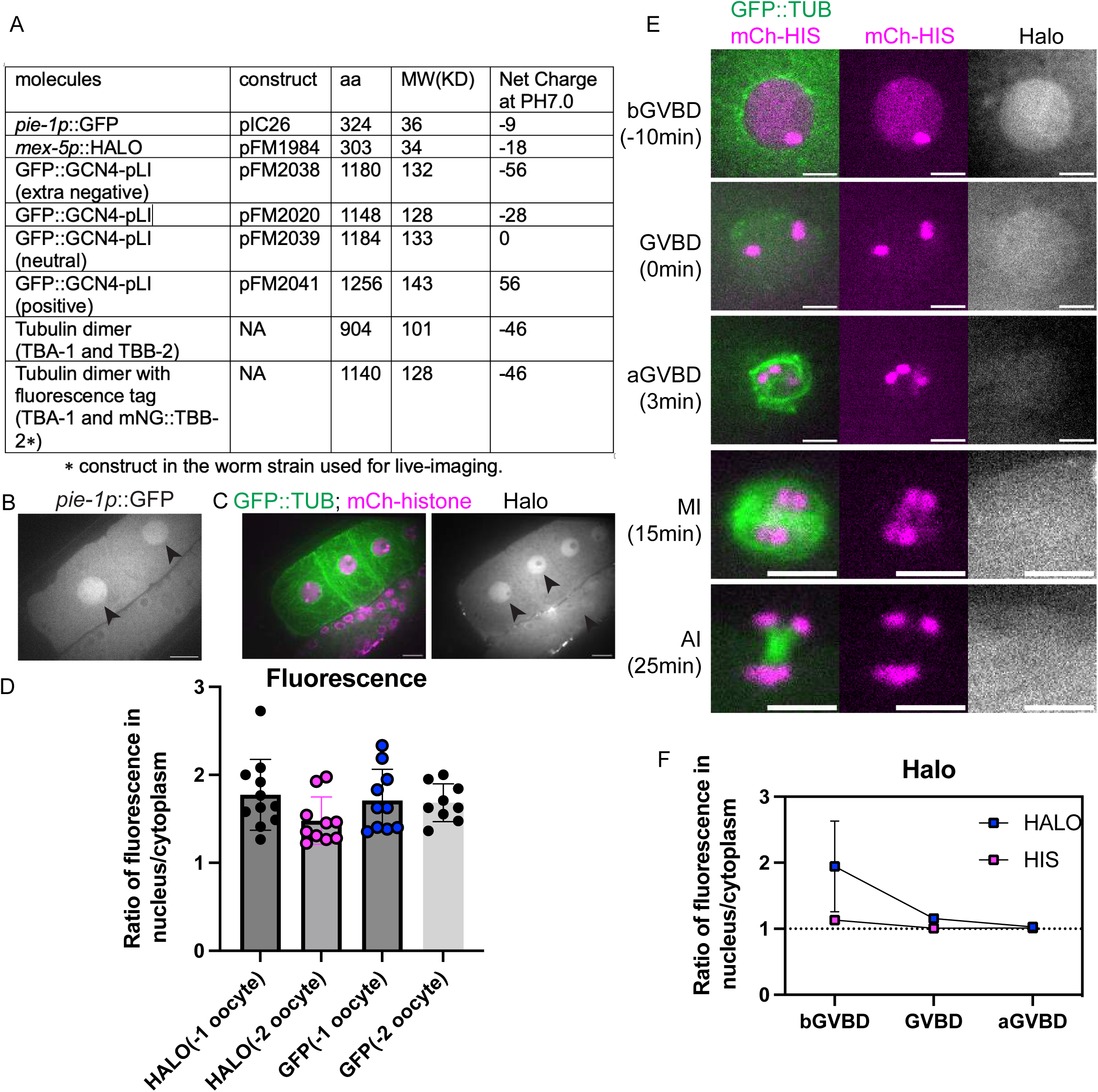
Molecule movement during GVBD is size dependent. (A) Molecular weight and net charge of molecules used in this study, expressed in the *C. elegans* germline. (B) Images of diakinesis oocytes expressing GFP (greyscale) before Germinal Vesicle Breakdown, and (C) Diakinesis oocytes expressing GFP::TUB (green), mCh::histone (magenta) and HALO (greyscale). Scale Bars, 10um. (D) Fluorescence intensity ratio of GFP or HALO in the nucleus to the cytoplasm in -1 or -2 oocytes. (E) Time lapse images of meiotic embryo expressing GFP::TUB (green), mCh::histone (magenta) and HALO (greyscale). Scale Bars, 5um. (F) Fluorescence intensity ratio of HALO and non-chromosome histone in the nucleus to the cytoplasm before GVBD, at GVBD onset and after GVBD.

These results suggested that the same mechanisms driving concentration of larger proteins during GVBD might be responsible for the concentration of smaller proteins before GVBD. However, monomeric HALO tag did not stay concentrated after GVBD and instead diffused to a 1:1 fluorescence ratio inside and outside the spindle envelope (Figure 7E-F). The monomeric HALO tag did not grossly perturb meiotic progression (Figure 5E) in 12/12 time-lapse sequences. These results suggested that there might be a size-dependence for concentration of proteins in the nuclear volume during meiotic spindle assembly.

### Molecule movement during GVBD is charge dependent

Although the tubulin-sized molecules that concentrated in the nuclear volume at GVBD are larger than the smaller HALO tag that quickly dispersed to a 1:1 ratio between nuclear and cytoplasmic volumes at GVBD, these proteins also differ in net charge (Fig. 7A), with the concentrating proteins more negative than the dispersing proteins. Single molecule diffusion studies in cytoplasm (Xiang et al., 2020) and inside organelles (Xiang et al., 2023) have revealed that net positive charge can slow or stop diffusion of small proteins. One possibility is that proteins with positive charge would interact transiently by ion exchange with negatively charged phosphatidyl serine-rich intracellular membranes, this electrostatic interaction would slow diffusion of positively charged proteins into a fenestrated spindle envelope but would not affect diffusion of negatively charged proteins. To test this idea, we added arginines to GFP::GCN4-pLI to either neutralize net charge or add a net positive charge. Similar to GFP::GCN4-pLI (Figure 7A, net charge at pH 7.0: -28), GFP::GCN4-pLI (neutral) or GFP::GCN4-pLI (positive) (Figure 7A, net charge at pH 7.0: 0 or 56 respectively) were excluded from the nucleus before GVBD (Figure 8A-D; -10 min). GFP::GCN4-pLI (neutral) remained excluded from the nuclear volume at GVBD for a longer period of time after histone leakage out of the nucleus than—GFP::GCN4-pLI and exhibited only a slight accumulation in the nuclear volume (∼7min after GVBD onset) (Figure 8A, 8C), a significant delay compared to GFP::GCN4-pLI (∼2min after GVBD onset, Figure 4C). GFP::GCN4-pLI (positive) remained excluded from the nuclear volume for an even longer period after GVBD (Figure 8B, 8D). This result suggested that proteins with net negative charge concentrate in the nuclear volume at GVBD whereas proteins with neutral or positive charges do not. Interestingly, GFP::GCN4-pLI (extra negative) (Figure 7, net charge at PH 7.0: -56) did not accumulate in the nuclear volume as GFP::GCn4-pLI did (Figure S5), suggesting there might be a narrow range of charge that are not retained in the cytoplasm.

**Fig 8.**
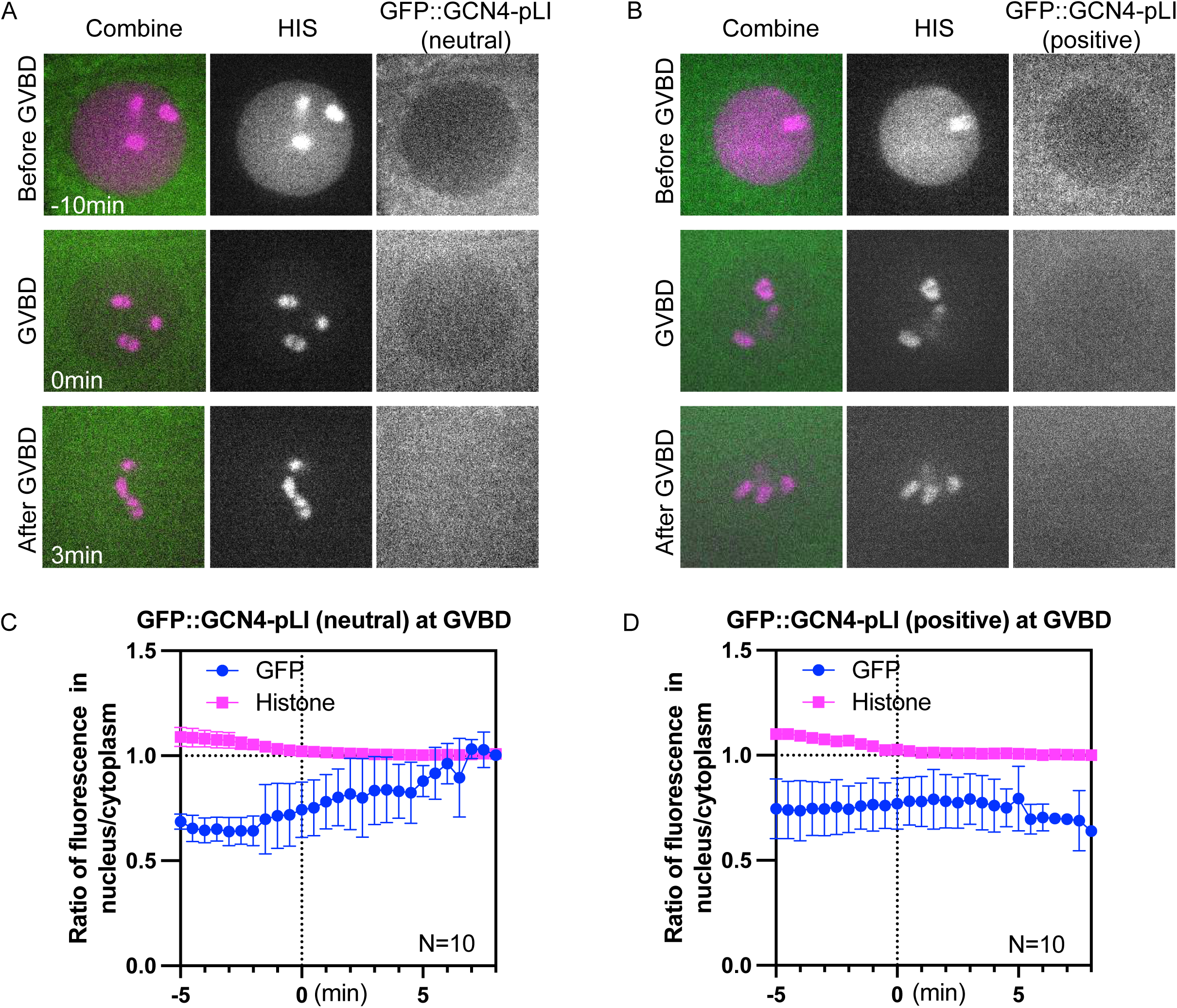
Molecule movement during GVBD is charge dependent. (A) Representative time lapse images of -1 oocyte expressing mCh::histone and GFP::GCN4-pLI (neutral) or (B) GFP::GCN4-pLI (positive) (C) and (D) Plots of fluorescence intensity ratio in nucleus to cytoplasm over time in (A) and (B), respectively. Y axis: fluorescence intensity [nucleus-background] ÷ fluorescence intensity [cytoplasm-background]. N: number of time lapse sequences analyzed. Mean is shown in solid magenta square [His] or solid green circle [GFP]. Bars indicate SEM.

## Discussion

Our data suggests that the ran pathway is dispensable for spindle formation but is important for limiting meiotic anaphase velocity and polar body extrusion. This is consistent with the localization of endogenously tagged Ran-GEF, RAN-3, which dispersed from chromosomes during meiotic spindle assembly but then concentrated on chromosomes during anaphase. It remains possible that the ran pathway is redundant with another pathway, that there is chromosome-associated RAN-3 during spindle assembly that is below our detection limit, or that if RAN-3 were more completely depleted, then an assembly defect would be observed. It is also possible that other chromatin-associated activators of spindle assembly, such as the CPC (Divekar et al., 2021), katanin (McNally et al., 2006; 2014), and CLS-2 (Schlientz and Bowerman, 2020) substitute for Ran-GTP during *C. elegans* meiosis more robustly than in other species.

Increased anaphase velocities have been reported after double depletions of three microtubule crosslinkers, KLP-19, BMK-1, and SPD-1 (Li et al., 2023). Thus Ran-GEF on anaphase chromosomes might generate Ran-GTP that would locally activate one or more microtubule crosslinkers in the anaphase spindle. RNAi depletion of RAN-3 and RAN-2 had the opposite effect, slowing anaphase, in mitotic spindles with ablated centrosomes (Nahaboo et al., 2015), suggesting different mechanisms at work during meiosis vs mitosis. Ran-GEF on anaphase chromosomes is closely juxtaposed against the cortex where Ran-GTP might activate components of the acto-myosin machinery to promote polar body formation as occurs in mouse oocytes (Deng et al., 2007).

The concentration of soluble tubulin in the nuclear volume during and after GVBD in *C. elegans* oocytes treated with nocodazole recapitulates what has been reported in mitotic cells of *C. elegans* and *Drosophila* (Hayashi et al., 2012; Métivier et al., 2021; Schweizer et al., 2015; Yao et al., 2012). Remaining questions are the mechanism and significance of this phenomenon. Hayashi et al., 2012 concluded that tubulin binds to something in the nuclear volume because of a fraction of nocodazole-treated GFP::tubulin that recovered slowly after photobleaching. Métivier et al., 2021 concluded that tubulin binds to tubulin folding cofactor E in the nucleus because depletion of cofactor E abrogated concentration of nocodazole-treated GFP::tubulin. Our finding that tetrameric GFP concentrates in the nuclear volume during GVBD supports the idea that a non-specific biophysical difference between the nucleoplasm and cytoplasm is responsible. This idea is consistent with the concentration of fluorescent dextrans (carbohydrates) in the spindle volume of *Drosophila* embryos (Yao et al., 2012) and the observation by Schweizer et al., 2015 that the mobility of nocodazole-treated GFP::tubulin molecules inside and outside the spindle volume are identical in S2 cells. Previous studies of nocodazole-treated GFP::tubulin could not address the cell cycle regulation of this phenomenon because nocodazole blocks spindle progression. We found that concentration of tetrameric GFP, or an unpolymerizable tubulin mutant during metaphase I and metaphase II correlated with the encasement of the spindle with layered sheets of ER and the dispersal during anaphase I and II correlated with dispersed tubular ER. The idea that a semipermeable envelope around the spindle is required for the concentration of soluble tubulin was supported by Schweizer et al., 2015 who showed that laser cutting of the spindle envelope prevents the concentration of soluble tubulin. Genetic or physical perturbation of the ER in *C. elegans* could help to address both the mechanism of tubulin concentration and the significance. We expect that any perturbation that disrupts the ER spindle envelope would prevent concentration of free tubulin and cause a delay in spindle assembly. If pathways are redundant, severe spindle assembly defects might only be observed after combining depletions with other chromosomal spindle assembly factors like RAN-3, katanin, or the CPC.

Our results indicate that both size and net charge determine whether a protein will concentrate in the nuclear volume before and after GVBD. We suggest that sheet-like ER is required to exclude mitochondria, yolk granules, and other organelles from the spindle volume. The cytoplasmic faces of cytoplasmic organelles are thought to be negatively charged due to phosphoinositides and phosphatidylserine. A recent study mapping the electrostatic profile of cellular membranes suggests that plasma membrane, ER, mitochondria, Golgi are all negatively charged in Hela cells (Eisenberg et al., 2021; average surface potential ranging from -14mV to -35mV). The negatively charged lipid heads will have counterions neutralizing these charges. However, it is possible the negative surface charge of cellular membranes might act like cation exchange chromatography beads, causing transient binding of proteins with net positive charges. In this scenario, diffusion of positively charged proteins into the spindle volume would be slowed, whereas diffusion out to the cytoplasm would be unrestricted. In contrast, proteins with a strong net negative charge would diffuse freely in both directions but appear concentrated in the spindle volume because of the low available volume between organelles in the cytoplasm. One logical outcome of this model is that tubulin concentration at the nanoscale between organelles would actually be no different than in the nuclear/spindle volume. However, the total mass of microtubules that could polymerize would be severely restricted, especially if a polymerizing plus end was stimulated to catastrophe by collision with an organelle. These ideas could readily be tested with in vitro experiments.

A recent study found that a katanin mutant that cannot bind microtubules still concentrates in the spindle volume (Beaumale et al., 2024) suggesting that other spindle assembly factors besides tubulin may be concentrated by the same biophysical mechanism.

## Materials and Methods

### C. elegans strains

C. elegans strains used in this study are listed in Table S1.

### Transgenes generated for this study

syb7781 and syb7819 were generated by inserting Auxin Induced Degron (AID) and HALO tag sequences to the endogenous *ran-3* and *ran-2* loci through CRISPR/Cas9-mediated genome editing by Suny Biotech. Sequences of these insertions are in Supplemental data 1. *duSi18, duSi20, duSi23, duSi24* and *duSi25* were generated by inserting GFP::GCN4-pLI (or its charge variants), or GFP::tba-2(T349E) via Flp Recombinase-Mediated Cassette Exchange (RMCE) method (Nonet, 2020). The plasmids containing corresponding DNAs [GFP::GCN4-pLI, GFP::GCN4-pLI (extra negative), GFP::GCN4-pLI (neutral), GFP::GCN4-pLI (positive), GFP::tba-2(T349E)] were injected into *jsSi1579* landing site in NM5402 by In Vivo Biosystems (Eugene, OR), then the NeonGreen + FLP transgene was removed by outcrossing. Plasmids for RMCE insertion are constructed as follows: DNA sequences of GCN4-pLI and tba-2(T349E) were obtained as gBlocks from IDT (Coralville, IA) followed by restriction digestion and cloned into the vector by T4 DNA ligation. The vector is designed for RMCE insertion, and it contains GFP optimized to reduce germline silencing. For charge variants of GFP::GCN4-pLI, gblocks containing Aspartic Acid and Glutamic Acid were cloned into GFP::GCN4-pLI to add negative net charge. To neutralize negative charge of GFP::GCN4-pLI or add positive net charge, 7 arginine or 21 arginine were cloned onto GFP::GCN4-pLI. The strains were sequenced, and the sequences are listed in Supplemental data 2.

### Drug treatment

The strain used for nocodazole experiments has a *bus-17(e2800)* mutation which makes the worms cuticle permeable to drugs. 5mg/ml stock nocodazole solution (Sigma-Aldrich, St. Louis, MO, dissolved in 100% DMSO) was diluted into tricaine/tetramisole anesthetics to 5ug/ml just before adding to worms for live imaging. 100% DMSO without nocodazole were diluted in the same way for control treatment.

### Live imaging and Statistical Analysis

Worms were anesthetized with tricaine/tetramisole as described (Kirby et al. 1990; McCarter et al. 1999) and gently mounted between a coverslip and a thin 2% agarose pad on a slide. All time lapse images were captured with a Solamere spinning disk confocal microscope equipped with an Olympus IX-70 stand, Yokogawa CSU10, either Hamamatsu ORCA FLASH 4.0 CMOS (complementary metal oxide semiconductor) detector or Hamamatsu ORCA-Quest qCMOS (quantitative complementary metal oxide semiconductor) detector, Olympus 100x UPlanApo 1.35 objective, 100-mW Coherent Obis laser set at 30% power, and MicroManager software control. Pixel size was 65nm for the ORCA FLASH 4.0 CMOS detector and 46nm for the ORCA-Quest qCMOS detector. For measuring fluorescence intensity or nucleus sizes (in Figure 2A-C; Figure 7B-C; Figure S1A), z-stack images (except in Fig S2A) were taken on the same microscope, with 1-micron step size to capture the center of nucleus for measurements. Images in Fig S3A were captured with a Zeiss Laser Scanning microscope (LSM) 980, Zeiss Objective LD LCI Plan-Apochromat 40x/1.2 Imm Corr DIC M27 for water, silicon oil or glycerine. For time-lapse movies of GVBD and meiosis, images were captured every 30s. The fluorescence intensity in the background (where there was no worm) were subtracted before the fluorescence in the nucleus or cytoplasm before quantification.

For segmentation of ER in electron tomograms, ER structures were manually assigned at 10nm intervals using IMOD software (https://bio3d.colorado.edu/imod/). Ribosomes in 1µm^2^ sections of spindle and cytoplasm in both metaphase I and metaphase II tomograms were counted using ImageJ software. Threshold values for each section were set to half of the peak value and the number of particles was determined. The average particle size in metaphase I spindles and cytoplasm, and metaphase II spindles and cytoplasm were: 30.9+/- 2.0nm, 28.0 +/- 1.2nm, 26.3 +/- 1.0nm, and 26.8 +/- 1.7nm respectively. Anaphase velocities were reported as the increase in distance between chromosome masses divided by time.

Area occupied by membranous organelles was determined from previously reported electron micrographs (Bembenek et al., 2007; Howe et al., 2001). Organelles were manually segmented using the Labkit plugin in Image J. Resulting segmented layers were exported as bitmap tifs and total area was determined with the Measure function in Image J.

Numerical values corresponding to data points in figures are included in Supplemental data 3.

## Acknowledgments

We thank Kent McDonald, Barbara Meyer and Josh Bembenek for sharing electron microscopy data and Shuyan Qui for generating the PLIN-1 strain. We thank Ho-yi Mak, Jim Priess, Arshad Desai, and the Caenorhabditis Genetics Center (CGC) for providing strains. The CGC is funded by the NIH Office of Research Infrastructure Programs (P40 OD010440).

This work was funded by National Institute of General Medical Sciences grant R35GM136241 to FJM and by the US Department of Agriculture/National Institute of Food and Agriculture Hatch project (1009162 to FJM).

## Author Contributions

T Gong conceptualization, formal analysis, validation, methodology, and writing—original draft

KL McNally formal analysis and methodology

S Konanoor formal analysis and methodology

A Peraza formal analysis and methodology

C Bailey formal analysis and methodology

S Redemann resources, formal analysis and methodology

FJ McNally conceptualization, formal analysis, supervision, funding acquisition, validation, project administration, and writing—original draft, review, and editing

## Figure Legends

**Fig S1. Ran-GEF and Ran-GAP are not required for meiotic spindle assembly.** (A) Images of RAN-3::AID::HALO, TIR1::mRuby, GFP::TUB and mCh::histone -1 oocytes before Germinal Vesicle Breakdown (bGVBD) and after (aGVBD). (B) Fluorescence intensity ratios of HALO in oocytes treated with 0, 4 or 6-hour Auxin. Control: strain not containing AID::HALO tag. (C) Images of RAN-2::AID::HALO, TIR1::mRuby, GFP::TUB and mCh::histone -1 oocytes before Germinal Vesicle Breakdown (bGVBD) and after (aGVBD). (D) Fluorescence intensity ratios of HALO in oocytes treated with 0, 4 or 6-hour Auxin. Control: strain not expressing AID::HALO. (E) Metaphase I spindle lengths determined from time-lapse. (F) Metaphase II spindle lengths determined from time-lapse. (G) Anaphase II velocities determined from time-lapse. ns P > 0.05, *P ≤ 0.05, Mann-Whitney U test. Size Bars, 5um.

**Fig S2. RAN-3::AID::HALO oocytes treated with auxin have pre-GVBD defects.**

(A) Representative images of oocytes expressing TIR1::mRuby, GFP::TUB (tubulin) and mCh::histone (mCherry) or RAN-3::AID::HALO, TIR1::mRuby, GFP::TUB and mCh::histone, treated with no auxin, 4hr auxin or 36hr auxin, respectively. Scale bars, 5μm.

(B) Quantification of oocyte nuclei size.

(C) Quantification of microtubule fluorescence ratio in nucleus to cytoplasm as an indication of nuclear envelope leakiness. ns P > 0.05, *P ≤ 0.05, **P ≤ 0.01, ***P ≤ 0.001, ****P ≤ 0.0001, Mann-Whitney U test. Size Bars, 5um.

**Fig S3. Difference of tubulin or GCN4-pLI fluorescence between nucleus and cytoplasm measured from Zeiss LSM confocal microscope is greater than from spinning disk confocal.**

(A) Representative images of diakinesis oocytes expressing mNG::TBB-2 (mNeonGreen:: tubulin) and mCh::HIS (mCherry::histone H2b) from Zeiss LSM confocal microscope. Scale bars, 10μm.

(B) Comparison of fluorescence intensity ratios of nucleus/cytoplasm in images acquired with a spinning disk confocal vs a laser scanning confocal.

(C) Representative images of oocytes expressing GFP::GCN4-pLI and mCh::HIS from Zeiss LSM confocal microscope. Scale bars, 10μm.

(D) Ratio of GFP::GCN4-pLI fluorescence in the nucleus/cytoplasm from images captured on a spinning disk confocal vs a laser scanning confocal.

**Fig. S4. Cytoplasmic volume determined from transmission electron micrograph.**

(A) High magnification of metaphase II spindle in the embryo on the right in B to show the quality of fixation.

(B) Low magnification TEM of -1 oocyte to metaphase I embryo.

(C) Manually segmented membrane vesicles of entire -1 oocyte and packed region outside the spindle in the meiotic embryo.

(D) Manually segmented non-vesicle regions.

**Fig S5. Movement of GFP::GCN4-pLI (extra negative) during GVBD.**

Plots of fluorescence intensity ratio in nucleus to cytoplasm over time in -1 oocytes expressing mCh::histone and GFP::GCN4-pLI (extra negative) during GVBD. Y axis: mean fluorescence intensity [nucleus-background] ÷ mean fluorescence intensity [cytoplasm-background]. N: number of time lapse sequences analyzed. Mean is shown in solid magenta square [His] or solid green circle [GFP].

**Video 1. Control meiosis I.** Control time-lapse sequence of meiosis I, 4 hr auxin treatment of a strain expressing GFP::tubulin (green), mCherry::histone (magenta), and TIR1.

**Video 2. RAN-3-depleted anaphase I through anaphase II.**

Time-lapse sequence of meiosis I and II, 4 hr auxin treatment of a RAN-3::AID::HALO strain expressing GFP::tubulin (green), mCherry::histone (magenta), and TIR1. One set of chromosomes from anaphase I merge into the metaphase II spindle.

**Video 3. GFP::GCN4-pLI concentrated in the nuclear volume at GVBD and metaphase I.** Time lapse sequences of -1 oocytes expressing GFP::GCN4-pLI (green), HALO::ER (magenta) and mCherry::histone (red). GFP::GCN4-pLI concentrates in the nuclear volume at GVBD, then disperses at anaphase I when the ER envelope around the spindle disperses.

**Video 4. GFP::TBA-2(T349E) concentrated in the nuclear volume at GVBD.**

Time lapse sequences of -1 oocyte expressing GFP::TBA-2(T349E) (green) and mCherry::histone (magenta).

**Video 5. GFP::TBA-2(T349E) metaphase I through anaphase II.**

Time lapse sequences of meiotic embryo expressing GFP::TBA-2(T349E) (green) and mCherry::histone (red). GFP::TBA-2(T349E) is concentrated in the spindle volume at metaphase I, disperses at anaphase I, re-concentrates at metaphase II, then disperses again at anaphase II.

**Video 6. Z-stack moving through an electron tomogram of a metaphase II spindle**. The number of z planes has been reduced 20 fold to reduce file size. The two lobes of the 6 univalent chromosomes are oriented down the pole to pole axis. ER sheets appear as white tubules in a single z plane but are contiguous when moving between z planes.

**Video 7. ER sheets envelop the sides of a metaphase II spindle.**

Rotating model of ER sheets on the partial exterior of a metaphase II meiotic spindle and spanning 1.2 um in the z, derived from an electron tomogram and corresponding to Fig. 6A. ER in green. Bi-lobed univalent chromosome in magenta. Cytoplasmic organelles in blue.

**Video 8. ER at a metaphase II spindle pole.** Rotating model of complex ER structure at one pole of a metaphase II spindle and spanning 1.2 um in z, derived from an electron tomogram and corresponding to Fig. 6D. ER in green. Two bi-lobed univalent chromosomes in magenta. Cytoplasmic organelles in blue.

**Video 9. ER around an anaphase I spindle.**

Rotating model of ER in a 0.6 um z-section of a late anaphase I spindle, derived from an electron tomogram and corresponding to Fig. 6. ER in green. Two segregating chromosome masses in magenta. Cytoplasmic organelles in blue.

**Video 10. ER delimits the concentration of GFP::GCN4-pLI during mitosis.**

Time-lapse sequences of an embryo expressing GFP::GCN4-pLI (green); mCherry::histone (red) and HALO::ER (magenta) from pronuclear meeting through the 4-cell stage. GFP fluorescence concentrated in the nuclear volume during GVBD, after GVBD and at metaphase I. The concentration is enclosed by ER during metaphase and disperses when ER disperses during anaphase.

## References

Angell RR. 1991. Predivision in human oocytes at meiosis I: a mechanism for trisomy formation in man. Hum Genet 86:383–387. doi:10.1007/BF00201839

Askjaer P, Galy V, Hannak E, Mattaj IW. 2002. Ran GTPase cycle and importins alpha and beta are essential for spindle formation and nuclear envelope assembly in living Caenorhabditis elegans embryos. Mol Biol Cell 13:4355–4370. doi:10.1091/MBC.E02-06-0346

Baumann C, Wang X, Yang L, Viveiros MM. 2017. Error-prone meiotic division and subfertility in mice with oocyte-conditional knockdown of pericentrin. J Cell Sci 130:1251–1262. doi:10.1242/JCS.196188

Beaumale E, Van Hove L, Pintard L, Joly N. 2024. Microtubule-binding domains in Katanin p80 subunit are essential for severing activity in C. elegans. J Cell Biol 223. doi:10.1083/JCB.202308023

Bembenek JN, Richie CT, Squirrell JM, Campbell JM, Eliceiri KW, Poteryaev D, Spang A, Golden A, White JG. 2007. Cortical granule exocytosis in C. elegans is regulated by cell cycle components including separase. Development 134:3837–3848. doi:10.1242/DEV.011361

Bennabi I, Terret ME, Verlhac MH. 2016. Meiotic spindle assembly and chromosome segregation in oocytes. J Cell Biol 215:611–619. doi:10.1083/JCB.201607062

Bilbao-Cortés D, Hetzer M, Längst G, Becker PB, Mattaj IW. 2002. Ran Binds to Chromatin by Two Distinct Mechanisms. Curr Biol 12:1151–1156. doi:10.1016/S0960-9822(02)00927-2

Bornens M. 2012. The centrosome in cells and organisms. Science (80-) 335:422–426. doi:10.1126/SCIENCE.1209037

Carazo-Salas RE, Guarguaglini G, Gruss OJ, Segref A, Karsenti E, Mattaj LW. 1999. Generation of GTP-bound Ran by RCC1 is required for chromatin-induced mitotic spindle formation. Nat 1999 4006740 400:178–181. doi:10.1038/22133

Cesario J, McKim KS. 2011. Ran-GTP is required for meiotic spindle organization and the initiation of embryonic development in Drosophila. J Cell Sci 124:3797–3810. doi:10.1242/JCS.084855

Chuang CH, Schlientz AJ, Yang J, Bowerman B. 2020. Microtubule assembly and pole coalescence: Early steps in Caenorhabditis elegans oocyte meiosis I spindle assembly. Biol Open 9. doi:10.1242/BIO.052308

Colombié N, Głuszek AA, Meireles AM, Ohkura H. 2013. Meiosis-Specific Stable Binding of Augmin to Acentrosomal Spindle Poles Promotes Biased Microtubule Assembly in Oocytes. PLOS Genet 9:e1003562. doi:10.1371/JOURNAL.PGEN.1003562

Conduit PT, Wainman A, Raff JW. 2015. Centrosome function and assembly in animal cells. Nat Rev Mol Cell Biol 2015 1610 16:611–624. doi:10.1038/nrm4062

Delarue M, Brittingham GP, Pfeffer S, Surovtsev I V., Pinglay S, Kennedy KJ, Schaffer M, Gutierrez JI, Sang D, Poterewicz G, et al. 2018. mTORC1 Controls Phase Separation and the Biophysical Properties of the Cytoplasm by Tuning Crowding. Cell 174:338–349.e20. doi:10.1016/J.CELL.2018.05.042

Deng M, Suraneni P, Schultz RM, Li R. 2007. The Ran GTPase mediates chromatin signaling to control cortical polarity during polar body extrusion in mouse oocytes. Dev Cell. 2007 Feb;12(2):301-8. doi: 10.1016/j.devcel.2006.11.008.

Divekar NS, Davis-Roca AC, Zhang L, Dernburg AF, Wignall SM. 2021. A degron-based strategy reveals new insights into Aurora B function in C. elegans. PLOS Genet 17:e1009567. doi:10.1371/JOURNAL.PGEN.1009567

Drutovic D, Duan X, Li R, Kalab P, Solc P. 2020. Ran GTP and importin β regulate meiosis I spindle assembly and function in mouse oocytes. EMBO J 39:e101689. doi:10.15252/embj.2019101689

Dumont J, Desai A. 2012. Acentrosomal spindle assembly and chromosome segregation during oocyte meiosis. Trends Cell Biol 22:241–249. doi:10.1016/j.tcb.2012.02.007

Dumont J, Petri S, Pellegrin F, Terret ME, Bohnsack MT, Rassinier P, Georget V, Kalab P, Gruss OJ, Verlhac MH. 2007. A centriole- and RanGTP-independent spindle assembly pathway in meiosis I of vertebrate oocytes. J Cell Biol 176:295–305. doi:10.1083/JCB.200605199

Eisenberg S, Haimov E, Walpole GFW, Plumb J, Kozlov MM, Grinstein S. 2021. Mapping the electrostatic profiles of cellular membranes. Mol Biol Cell 32:301–310. doi:10.1091/MBC.E19-08-0436

Fair T, Lonergan P. 2023. The oocyte: the key player in the success of assisted reproduction technologies. Reprod Fertil Dev. doi:10.1071/RD23164

Frasch M. 1991. The maternally expressed Drosophila gene encoding the chromatin-binding protein BJ1 is a homolog of the vertebrate gene Regulator of Chromatin Condensation, RCC1. EMBO J 10:1225–1236. doi:10.1002/J.1460-2075.1991.TB08064.X

Galy V, Mattaj IW, Askjaer P. 2003. Caenorhabditis elegans Nucleoporins Nup93 and Nup205 Determine the Limit of Nuclear Pore Complex Size Exclusion In Vivo. Mol Biol Cell 14:5104. doi:10.1091/MBC.E03-04-0237

Goshima G, Mayer M, Zhang N, Stuurman N, Vale RD. 2008. Augmin: A protein complex required for centrosome-independent microtubule generation within the spindle. J Cell Biol 181:421–429. doi:10.1083/JCB.200711053

Gruss OJ. 2018. Animal Female Meiosis: The Challenges of Eliminating Centrosomes. Cells 2018, Vol 7, Page 73 7:73. doi:10.3390/CELLS7070073

Halpin D, Kalab P, Wang J, Weis K, Heald R. 2011. Mitotic Spindle Assembly around RCC1-Coated Beads in Xenopus Egg Extracts. PLOS Biol 9:e1001225. doi:10.1371/JOURNAL.PBIO.1001225

Hayashi H, Kimura K, Kimura A. 2012. Localized accumulation of tubulin during semi-open mitosis in the Caenorhabditis elegans embryo. Mol Biol Cell 23:1688–1699. doi:10.1091/MBC.E11-09-0815

Heald R, Tournebize R, Habermann A, Karsenti E, Hyman A. 1997. Spindle Assembly in Xenopus Egg Extracts: Respective Roles of Centrosomes and Microtubule Self-Organization. J Cell Biol 138:615–628. doi:10.1083/JCB.138.3.615

Hetzer M, Gruss OJ, Mattaj IW. 2002. The Ran GTPase as a marker of chromosome position in spindle formation and nuclear envelope assembly. Nat Cell Biol 2002 47 4:E177–E184. doi:10.1038/ncb0702-e177

Hinchcliffe EH. 2014. Centrosomes and the Art of Mitotic Spindle Maintenance. Int Rev Cell Mol Biol 313:179–217. doi:10.1016/B978-0-12-800177-6.00006-2

Holubcová Z, Blayney M, Elder K, Schuh M. 2015. Error-prone chromosome-mediated spindle assembly favors chromosome segregation defects in human oocytes. Science (80-) 348:1143–1147. doi:10.1126/SCIENCE.AAA9529

Howe M, McDonald KL, Albertson DG, Meyer BJ. 2001. Him-10 Is Required for Kinetochore Structure and Function on Caenorhabditis elegans Holocentric Chromosomes. J Cell Biol 153:1227–1238. doi:10.1083/JCB.153.6.1227

Johnson V, Ayaz P, Huddleston P, Rice LM. 2011. Design, Overexpression, and Purification of Polymerization-Blocked Yeast αβ-Tubulin Mutants. Biochemistry 50:8636–8644. doi:10.1021/BI2005174

Kalab P, Pu RT, Dasso M. 1999. The Ran GTPase regulates mitotic spindle assembly. Curr Biol 9:481–484. doi:10.1016/S0960-9822(99)80213-9

Kellogg DR, Moritz M, Alberts BM. 2003. THE CENTROSOME AND CELLULAR ORGANIZATION. https://doi.org/101146/annurev.bi63070194003231 63:639–674. doi:10.1146/ANNUREV.BI.63.070194.003231

Khodjakov A, Cole RW, Oakley BR, Rieder CL. 2000. Centrosome-independent mitotic spindle formation in vertebrates. Curr Biol 10:59–67. doi:10.1016/S0960-9822(99)00276-6

Kline-Smith SL, Walczak CE. 2004. Mitotic Spindle Assembly and Chromosome Segregation: Refocusing on Microtubule Dynamics. Mol Cell 15:317–327. doi:10.1016/J.MOLCEL.2004.07.012

Kraus J, Travis SM, King MR, Petry S. 2023. Augmin is a Ran-regulated spindle assembly factor. J Biol Chem 299. doi:10.1016/j.jbc.2023.104736

Lantzsch I, Yu CH, Chen YZ, Zimyanin V, Yazdkhasti H, Lindow N, Szentgyoergyi E, Pani AM, Prohaska S, Srayko M, et al. 2021. Microtubule reorganization during female meiosis in c. Elegans. Elife 10. doi:10.7554/ELIFE.58903

Lawo S, Bashkurov M, Mullin M, Ferreria MG, Kittler R, Habermann B, Tagliaferro A, Poser I, Hutchins JRA, Hegemann B, et al. 2009. HAUS, the 8-subunit human Augmin complex, regulates centrosome and spindle integrity. Curr Biol 19:816–826. doi:10.1016/J.CUB.2009.04.033

Lénárt P, Rabut G, Daigle N, Hand AR, Terasaki M, Ellenberg J. 2003. Nuclear envelope breakdown in starfish oocytes proceeds by partial NPC disassembly followed by a rapidly spreading fenestration of nuclear membranes. J Cell Biol 160:1055–1068. doi:10.1083/jcb.200211076

Li W, Crellin HA, Cheerambathur D, McNally FJ. 2023. Redundant microtubule crosslinkers prevent meiotic spindle bending to ensure diploid offspring in *C. elegans*. PLos Genetics. 19(12):e1011090.

Li HY, Wirtz D, Zheng Y. 2003. A mechanism of coupling RCC1 mobility to RanGTP production on the chromatin in vivo. J Cell Biol 160:635. doi:10.1083/JCB.200211004

Li X, Qin Y, Wilsher S, Allen WR. 2006. Centrosome changes during meiosis in horse oocytes and first embryonic cell cycle organization following parthenogenesis, fertilization and nuclear transfer. Reproduction 131:661–667. doi:10.1530/REP.1.00795

McNally K, Audhya A, Oegema K, McNally FJ. 2006. Katanin controls mitotic and meiotic spindle length. J Cell Biol 175:881–891. doi:10.1083/jcb.200608117

McNally K, Berg E, Cortes DB, Hernandez V, Mains PE, McNally FJ. 2014. Katanin maintains meiotic metaphase chromosome alignment and spindle structure in vivo and has multiple effects on microtubules in vitro. Mol Biol Cell 25:1037–1049. doi:10.1091/mbc.E13-12-0764

McNally KP, Beath EA, Danlasky BM, Barroso C, Gong T, Li W, Martinez-Perez E, McNally FJ. 2022. Cohesin is required for meiotic spindle assembly independent of its role in cohesion in C. elegans. PLOS Genet 18:e1010136. doi:10.1371/JOURNAL.PGEN.1010136

Métivier M, Gallaud E, Thomas A, Pascal A, Gagné JP, Poirier GG, Chrétien D, Gibeaux R, Richard-Parpaillon L, Benaud C, et al. 2021. Drosophila Tubulin-Specific Chaperone E Recruits Tubulin around Chromatin to Promote Mitotic Spindle Assembly. Curr Biol 31:684–695.e6. doi:10.1016/J.CUB.2020.11.009

Mikeladze-Dvali T, von Tobel L, Strnad P, Knott G, Leonhardt H, Schermelleh L, Gönczy P. 2012. Analysis of centriole elimination during C. elegans oogenesis. Development 139:1670–1679. doi:10.1242/DEV.075440

Mittl PRE, Deillon C, Sargent D, Liu N, Klauser S, Thomas RM, Gutte B, Grütter MG. 2000. The retro-GCN4 leucine zipper sequence forms a stable three-dimensional structure. Proc Natl Acad Sci U S A 97:2562–2566. doi:10.1073/PNAS.97.6.2562

Moore WJ, Zhang C, Clarke PR. 2002. Targeting of RCC1 to Chromosomes Is Required for Proper Mitotic Spindle Assembly in Human Cells. Curr Biol 12:1442–1447. doi:10.1016/S0960-9822(02)01076-X

Mullen TJ, Davis-Roca AC, Wignall SM. 2019. Spindle assembly and chromosome dynamics during oocyte meiosis. Curr Opin Cell Biol 60:53–59. doi:10.1016/J.CEB.2019.03.014

Mullen TJ, Wignall SM. 2017. Interplay between microtubule bundling and sorting factors ensures acentriolar spindle stability during C. elegans oocyte meiosis. PLoS Genet 13. doi:10.1371/journal.pgen.1006986

Nahaboo W, Zouak M, Askjaer P, Delattre M. 2015. Chromatids segregate without centrosomes during Caenorhabditis elegans mitosis in a Ran- and CLASP-dependent manner. Mol Biol Cell. 26(11):2020–9. doi: 10.1091/mbc.E14-12-1577. Epub 2015 Apr 1.

Nasmyth K. 2002. Segregating sister genomes: The molecular biology of chromosome separation. Science (80-) 297:559–565. doi:10.1126/science.1074757

Nguyen AL, Drutovic D, Vazquez BN, El Yakoubi W, Gentilello AS, Malumbres M, Solc P, Schindler K. 2018. Genetic Interactions between the Aurora Kinases Reveal New Requirements for AURKB and AURKC during Oocyte Meiosis. Curr Biol 28:3458–3468.e5. doi:10.1016/j.cub.2018.08.052

Ohtsubo M, Okazaki H, Nishimoto T. 1989. The RCC1 protein, a regulator for the onset of chromosome condensation locates in the nucleus and binds to DNA. J Cell Biol 109:1389–1397. doi:10.1083/JCB.109.4.1389

Olson SK, Greenan G, Desai A, Müller-Reichert T, Oegema K. 2012. Hierarchical assembly of the eggshell and permeability barrier in C. elegans. J Cell Biol. 2012 Aug 20;198(4):731-48. doi: 10.1083/jcb.201206008

Penfield L, Shankar R, Szentgyörgyi E, Laffitte A, Mauro MS, Audhya A, Müller-Reichert T, and Bahmanyar S. 2020. Regulated lipid synthesis and LEM2/CHMP7 jointly control nuclear envelope closure. J Cell Biol. 2020 May 4; 219(5): e201908179.

Petry S. 2016. Mechanisms of Mitotic Spindle Assembly. https://doi.org/101146/annurev-biochem-060815-014528 85:659–683. doi:10.1146/ANNUREV-BIOCHEM-060815-014528

Petry S, Groen AC, Ishihara K, Mitchison TJ, Vale RD. 2013. Branching microtubule nucleation in xenopus egg extracts mediated by augmin and TPX2. Cell 152:768–777. doi:10.1016/j.cell.2012.12.044

Petry S, Pugieux C, Nedeĺec FJ, Vale RD. 2011. Augmin promotes meiotic spindle formation and bipolarity in Xenopus egg extracts. Proc Natl Acad Sci U S A 108:14473–14478. doi:10.1073/PNAS.1110412108

Poteryaev D, Squirrell JM, Campbell JM, White JG, Spang A. 2005. Involvement of the actin cytoskeleton and homotypic membrane fusion in ER dynamics in Caenorhabditis elegans. Mol Biol Cell 16:2139–2153. doi:10.1091/MBC.E04-08-0726

Prosser SL, Pelletier L. 2017. Mitotic spindle assembly in animal cells: a fine balancing act. Nat Rev Mol Cell Biol 2017 183 18:187–201. doi:10.1038/nrm.2016.162

Sánchez-Huertas C, Lüders J. 2015. The Augmin Connection in the Geometry of Microtubule Networks. Curr Biol 25:R294–R299. doi:10.1016/J.CUB.2015.02.006

Sanchez AD, Feldman JL. 2017. Microtubule-organizing centers: from the centrosome to non-centrosomal sites. Curr Opin Cell Biol 44:93–101. doi:10.1016/J.CEB.2016.09.003

Schuh M, Ellenberg J. 2007. Self-Organization of MTOCs Replaces Centrosome Function during Acentrosomal Spindle Assembly in Live Mouse Oocytes. Cell 130:484–498. doi:10.1016/J.CELL.2007.06.025

Schlientz AJ, Bowerman B. 2020. *C. elegans* CLASP/CLS-2 negatively regulates membrane ingression throughout the oocyte cortex and is required for polar body extrusion. PLoS Genet. 16(10):e1008751. doi: 10.1371/journal.pgen.1008751.

Schwarzerová K, Bellinvia E, Martinek J, Sikorová L, Dostál V, Libusová L, Bokvaj P, Fischer L, Schmit AC, Nick P. 2019. Tubulin is actively exported from the nucleus through the Exportin1/CRM1 pathway. Sci Rep. 9(1):5725. doi: 10.1038/s41598-019-42056-6.

Schweizer N, Pawar N, Weiss M, Maiato H. 2015. An organelle-exclusion envelope assists mitosis and underlies distinct molecular crowding in the spindle region. J Cell Biol 210:695–704. doi:10.1083/jcb.201506107

So C, Menelaou K, Uraji J, Harasimov K, Steyer AM, Seres KB, Bucevičius J, Lukinavičius G, Möbius W, Sibold C, et al. 2022. Mechanism of spindle pole organization and instability in human oocytes. Science (80-) 375. doi:10.1126/SCIENCE.ABJ3944

Srayko M, Buster DW, Bazirgan OA, McNally FJ, Mains PE. 2000. MEI-1/MEI-2 katanin-like microtubule severing activity is required for Caenorhabditis elegans meiosis. Genes Dev 14:1072–84.

Thomas C, Cavazza T, Schuh M. 2021. Aneuploidy in human eggs: contributions of the meiotic spindle. Biochem Soc Trans 49:107–118. doi:10.1042/BST20200043

Uehara R, Nozawa RS, Tomioka A, Petry S, Vale RD, Obuse C, Goshima G. 2009. The augmin complex plays a critical role in spindle microtubule generation for mitotic progression and cytokinesis in human cells. Proc Natl Acad Sci U S A 106:6998–7003. doi:10.1073/PNAS.0901587106

Wang G, Jiang Q, Zhang C. 2014. The role of mitotic kinases in coupling the centrosome cycle with the assembly of the mitotic spindle. J Cell Sci 127:4111–4122. doi:10.1242/JCS.151753

Wolff ID, Tran M V., Mullen TJ, Villeneuve AM, Wignalla SM. 2016. Assembly of Caenorhabditis elegans acentrosomal spindles occurs without evident microtubuleorganizing centers and requires microtubule sorting by KLP-18/kinesin-12 and MESP-1. Mol Biol Cell 27:3122–3131. doi:10.1091/MBC.E16-05-0291

Wu T, Dong J, Fu J, Kuang Y, Chen B, Gu H, Luo Y, Gu R, Zhang M, Li W, et al. 2022. The mechanism of acentrosomal spindle assembly in human oocytes. Science (80-) 378. doi:10.1126/SCIENCE.ABQ7361

Xiang L, Chen K, Yan R, Li W, Xu K. 2020. Single-molecule displacement mapping unveils nanoscale heterogeneities in intracellular diffusivity. Nat Methods 2020 175 17:524–530. doi:10.1038/s41592-020-0793-0

Xiang L, Yan R, Chen K, Li W, Xu K. 2023. Single-Molecule Displacement Mapping Unveils Sign-Asymmetric Protein Charge Effects on Intraorganellar Diffusion. Nano Lett 23:1711–1716. doi:10.1021/ACS.NANOLETT.2C04379

Yao C, Rath U, Maiato H, Sharp D, Girton J, Johansen KM, Johansen J. 2012. A nuclear-derived proteinaceous matrix embeds the microtubule spindle apparatus during mitosis. Mol Biol Cell 23:3532–3541. doi:10.1091/MBC.E12-06-0429

Zhang L, Ward JD, Cheng Z, Dernburg AF. 2015. The auxin-inducible degradation (AID) system enables versatile conditional protein depletion in C. elegans. Dev 142:4374–4384. doi:10.1242/DEV.129635

