## Supplemental Data 1-2 and Table S1 for "Roles of Tubulin Concentration during Prometaphase and Ran-GTP during Anaphase of *C. elegans* meiosis"

**Supplemental Data 1: Sequences of ran degrons:**

ran-3(syb7781) II:
GGAAAAGTGTTTGCTATGGGAAAGAACACAGACAATGCTCTCGGCCTCGGTAATTG
GACTGGAAAGGACGACCAACAGCATTGGTTGTACGATACAATCCAGGAAATAGAATT
CGATTCGAAGATCGTTGGTGTTTCTGCCAAACTAGCCACTTCTATCGCCTGGTCTGA
GGATGGAACCGCCTACGCTTGGGGTTTTGATACTACCGGACAACTTGGTCTCGGAT
TGAAAGACGAAGACGAGAAGgtaatttttcaaaactcaaactatcaataaataaaatattcaatcttattttccag
ATGGTGTCCAAGCCAGAGGAGATCAGCTCC**GC**A**CA**CCTTGACGGTTATAGTATTAT
CGGGGCTTCGATTTCCGATCAGCACACTTTGATTATTGCCAAGAAAAATGGAGCATC
GGGAGCCTCAGGAGCATCGATGCCTAAAGATCCAGCCAAACCTCCGGCCAAGGCA
CAAGTTGTGGGATGGCCACCGGTGAGATCATACCGGAAGAACGTGATGGTTTCCTG
CCAAAAATCAAGCGGTGGCCCGGAGGCGGCGGCGTTCGTGAAGGGAGCATCGGG
AGCCTCAGGAGCATCGATGGCTGAAATTGGCACAGGATTCCCGTTTGACCCCCACT
ACGTCGAGGTCCTCGGAGAGCGTATGCACTACGTCGACGTCGGACCACGTGACGG
AACCCCAGTCCTCTTCCTCCACGGAAACCCAACCTCCTCCTACGTCTGGCGTAACA
TCATCCCACACGTCGCCCCAACCCACCGTTGCATCGCCCCAGACCTCATCGGAATG
GGAAAGTCCGACAAGCCAGACCTCGGATACTTCTTCGACGACCACGTCCGTTTCAT
GGACGCCTTCATCGAGGCCCTCGGACTCGAGGAGGTCGTCCTCGTCATCCACGAC
TGGGGATCCGCCCTCGGATTCCACTGGGCCAAGCGTAACCCAGAGCGTGTCAAGgt
aagtttaaacatatatatactaactaaccctgattatttaaattttcagGGAATCGCCTTCATGGAGTTCATCC
GTCCAATCCCAACCTGGGACGAGTGGCCAGAGTTCGCCCGTGAGACCTTCCAAGC
CTTCCGTACCACCGACGTCGGACGTAAGCTCATCATCGACCAAAACGTCTTCATCG
AGGGAACCCTCCCAATGGGAGTCGTCCGTCCACTCACCGAGGTCGAGATGGACCA
CTACCGTGAGCCATTCCTCAACCCAGTCGACCGTGAGCCACTCTGGCGTTTCCCAA
ACGAGCTCCCAATCGCCGGAGAGCCAGCCAACATCGTCGCCCTCGTCGAGGAGTA
CATGGACTGGCTCCACCAATCCCCAGTCCCAAAGCTCCTCTTCTGGGGAACCCCA
GGAGTCCTCATCCCACCAGCCGAGGCCGCCCGTCTCGCCAAGTCCCTCCCAAACT
GCAAGgtaagtttaaacagttcggtactaactaaccatacatatttaaattttcagGCCGTCGACATCGGACC
AGGACTCAACCTCCTCCAAGAGGACAACCCAGACCTCATCGGATCCGAGATCGCC
CGTTGGCTCTCCACCCTCGAGATCTCCGGATAAattatttggttttattctcaacttttatatcagttttgtttt
gtctctgtagcattattttgttattttttcgtttcccggtagccaattcgattgttctccagtaacattctcatcaatttttctgtttttttatc
atttcatttttgtcaagtagcatcagtcatgtaagaaagggatagagttcccttctgtgaaaatggagaattgttgaaacgtcgt
tgcacaacgacgtttaaacacttcactctatccctggattgatttccaacttctatgttttctcaaacccccctattagttctgctat
atattggaatccaaaaatttttcatttttagcctgagttatgttgttctcttatcatgtgaactcactgttttataatccgtttcaccattt
atggtaaacgtttcctgg

3’ of ran-3 are highlighted in yellow (within which synonymous mutation is labeled in blue text); AID in red text followed by halo sequence in blue text. Linker sequences (in purple) are inserted among the genes.

ran-2(syb7819) III:
AGCTATTGAAGTTGCAGgtaagaaatataaaaatattttaatataactactttcaatttttaagAAAATATCGT
CCGCCGAGTGGAGTCTGTCAAGCGTAACCCGATTCCGGCCACAACTCAATTAGTTA
ACAATATTGTTGCTCAATGTGCAGGAACAGGAGTTAAGgtaggtatttttcaagcttattctaaaaaa
cgtttaatatatgagacagactttacagGCTGAAACTGATTGGGGATATGGTGCCGATCCACAAG
TGATTTCACGTTTGTTCTCGGAACTTGTTGCTCGCGGCCATTTCAAGCTTGAGCTG
GCTCTCCTTCAACGCTTTTTCgtaagtctcacaactatattttatggttattttttcaatttttcagCCTTCACA
AGGAGCATCGGGAGCCTCAGGAGCATCGATGCCTAAAGATCCAGCCAAACCTCCG
GCCAAGGCACAAGTTGTGGGATGGCCACCGGTGAGATCATACCGGAAGAACGTGA
TGGTTTCCTGCCAAAAATCAAGCGGTGGCCCGGAGGCGGCGGCGTTCGTGAAGG
GAGCATCGGGAGCCTCAGGAGCATCGATGGCTGAAATTGGCACAGGATTCCCGTT
TGACCCCCACTACGTCGAGGTCCTCGGAGAGCGTATGCACTACGTCGACGTCGGA
CCACGTGACGGAACCCCAGTCCTCTTCCTCCACGGAAACCCAACCTCCTCCTACG
TCTGGCGTAACATCATCCCACACGTCGCCCCAACCCACCGTTGCATCGCCCCAGAC
CTCATCGGAATGGGAAAGTCCGACAAGCCAGACCTCGGATACTTCTTCGACGACCA
CGTCCGTTTCATGGACGCCTTCATCGAGGCCCTCGGACTCGAGGAGGTCGTCCTC
GTCATCCACGACTGGGGATCCGCCCTCGGATTCCACTGGGCCAAGCGTAACCCAG
AGCGTGTCAAGgtaagtttaaacatatatatactaactaaccctgattatttaaattttcagGGAATCGCCTTC
ATGGAGTTCATCCGTCCAATCCCAACCTGGGACGAGTGGCCAGAGTTCGCCCGTG
AGACCTTCCAAGCCTTCCGTACCACCGACGTCGGACGTAAGCTCATCATCGACCAA
AACGTCTTCATCGAGGGAACCCTCCCAATGGGAGTCGTCCGTCCACTCACCGAGG
TCGAGATGGACCACTACCGTGAGCCATTCCTCAACCCAGTCGACCGTGAGCCACT
CTGGCGTTTCCCAAACGAGCTCCCAATCGCCGGAGAGCCAGCCAACATCGTCGCC
CTCGTCGAGGAGTACATGGACTGGCTCCACCAATCCCCAGTCCCAAAGCTCCTCTT
CTGGGGAACCCCAGGAGTCCTCATCCCACCAGCCGAGGCCGCCCGTCTCGCCAA
GTCCCTCCCAAACTGCAAGgtaagtttaaacagttcggtactaactaaccatacatatttaaattttcagGCC
GTCGACATCGGACCAGGACTCAACCTCCTCCAAGAGGACAACCCAGACCTCATCG
GATCCGAGATCGCCCGTTGGCTCTCCACCCTCGAGATCTCCGGATAAtcgagcttctctac
acgatatcccagtcttcgcgatttttttacatgatttcagataagccgtggttattttatatttggatctacaatacatgtattattcat
cgtatcgtgaacaatatatttcttcaacttccaatgtatacacgagttatcaatttgttttaattttgtttctctaatttccactaattttta
gtttaatactttaaatctctttcttgtacgtgtaaatgcttcaatccgtttttcaagtaaatttttgtgaacgaagttgtttttatgattac
atgttttatgtctttgtgaactttttt

3’ of ran-2 are highlighted in yellow (within which synonymous mutation is labeled in blue text);  AID in red text followed by halo sequence in blue text. Linker sequences (in purple) are inserted among the genes.

**Supplemental Data 2: Sequences of transgene insertions:**

1. Sequence of GFP::GCN4-pLi:

ATGCCT**GCATcc**TCCAAGGGAGAGGAGCTCTTCACCGGAGTCGTCCCAATCCTCGTCGAGCTCGACGGAGgtattttcctgcatttttcaact
gggaaaatgaaagaaaatcgataatttcagACGTCAACGGACACAAGTTCTCCGTCTCCGGAGAGGGAGAGGGAGACGCCACCTACGGAAA
GCTCACCCTCAAGTTCATCTGCACCACCGGAAAGCTCCCAGTCCCATGGCCAACCCTCGTCACCACCTTCTGCTACGGAGgtaag
atatgggaaagaaggaaaaaaccgagattttacttgaaaaattgaatttttcgcgggattttcaccaaaaattgttgaatattcattatttcacgctgtaaaacaaaaaaaaaaaaaatcaaa
aactacgttgaaatcgcgtttttaagcgaattttcttcagaattgccagattttaaccccaaattttgcagtttttaaataaaatttcaccttttcggctcaaattgtagattttcctgaaaatttagtacaa
aaaacaatttcctcgtaaatttttcaaattttcagTCCAATGCTTCTCCCGTTACCCAGACCACATGAAGCGTCACGACTTCTTCAAGTCCGCCATG
CCAGAGGGATACGTCCAAGAGCGTACCATCTTCTTCAAGGACGACGGAAACTACAAGACCCGTGCCGAGGTCAAGTTCGAGGG
AGACACCCTCGTCAACCGTATCGAGCTCAAGGGAATCGACTTCAAGGAGGACGGAAACATCCTCGGACACAAGCTCGAGTACA
ACTACAACTCCCACAACGTCTACATCATGGCCGACAAGCAAAAGAACGGAATCAAGGTCAACTTCAAGgtacggattgaaattgctttaaaatt
tgaaaaattgattaaaaagtgcatttttaagctttgaccgacttaaaattagatttctgagcctattttctgagaattggaattttttcattgtgaaagttcaaagaatagcgtgaatgattagaaaa
tattgtaaaatttcaatttttcctataaaaaaggatttttttaggaatcaaaaattgcaaaatgatgccctaaaattcgaaaataaatataaaaattggcggtttctcaaaaatctagaattccgac
cttaattattaatttttttacaatattttttttgaaaaaaatccagaaaatttgaatttcgtagtttgtagtctaggcctctatcaattaaattttcgatttttgagtaaaatttcgaatttactattatttggacc
aaaattgtatttttttttcagaattaaatttataaaattttcagaaattaaatttataaaaaaaaaaaaaaaattaaaaaaaaaaaaaaaaaattaaaaattaaaaattaaaaaattaagtttttttta
ctcaaaattttgcactgaaattcgaaaatctaaaaatccgacctaaagtctgtattttttcaacaaaaattcagaaaaaactcaaaaacttgtattttgtagccagtcaaccactttctaaaatatc
aaatttgaatttttcaagcatttttacattgaaaaatctaatttttcgagtgaaaccacttgaaaatctgaaaattgaattaatttctgaactttagagatttttgtcaaaattttagataaagggattttt
aaacaaaaaattgatttttaatcgaaaaatactgggatttatgggtttttaaaagaaaaacgggggtttgaatgaaaaatcgcctgaaatcttagaaaattaattaaaaaactatgatttaatt
ccaaaaattaccgaaaatatcaaattttccatttttaaaccctaaattctttcagATCCGTCACAACATCGAGGACGGATCCGTCCAACTCGCCGACCACTAC
CAACAAAACACCCCAATCGGAGACGGACCAGTCCTCCTCCCAGACAACCACTACCTCTCCACCCAATCCGCCCTCTCCAAGgtag
attttttagaatttttgggtttttgaagtagaaaatcataaaaatctagggtttttatgaattgttttgaagaaaaattgcaaaaattccacaaaatggaagaaaaataactttggaagcgcatttttc
gcaaaaaaaccgaaatttttgcgtaaaatttcaaaattgcaataaaaattccacaaacatcaaatttcttaaatttttttataaaaaattggatggaaacactcttgaatttagaaaaaaaatcag
tttttctcatctaaaacttcaaaatttcggtgtaatcccattaaaattgccacaaaattcggaaatttcacctgaaatagagtgaataatttaaaatgttcagaaattcatattttgtcattttaaaagc
attaaaacaaatcaaaaatctattttttttggttggaaaagttcaaaattctggagaaattacattaaaattttcgtaaaatatggtaaataaacgaaaatgttgagaaattaaagaaaagttaca
atttttagcttaaaaattcaacattttgaggaaatgccacctaaaaaagtgactaatcgaaaatgttgagaaattaaaattgccaccatttatttatataaactactctaaaattacaattttcatgtt
aaaaattaataaaaaatctactttttccaaactacagtaaccttaccgtatacctacagtacctgaacattgccccccaccagctcccaacccaatacctcctcaaaaacttacacctcaatttt
tcataaactacagtaaccctaccaaaaaagcacaaaaaaaaaatctacattcattttccaacaattttcaatattttcagGACCCAAACGAGAAGCGTGACCACATGGTC
CTCCTCGAGTTCGTCACCGCCGCCGGAATCACCCACGGAATGGACGAGCTCTACAAGCCTGCATCaGGAGGcTCCGGtGGgTCg
GGtGGcTCaGGgGGAATGAAACAGATAGAGGATAAGCTTGAGGAGATTCTGTCCAAGCTGTATCACATTGAGAATGAGCTCGCTC
GCATCAAGAAGCTCTTGGGGGAGCGGTAA

GFP optimized for germline expression is highlighted in green. GCN4-pLi is colored red.

2. Sequence of GFP::GCN4-pLi with negative charge:

**"GAAGATGAAGATGAGGACGAGGCA"** was inserted onto GFP::GCN4-pLi before GCATcc (highlighted in bold)

3. Sequence of GFP::GCN4-pLi with neutral charge:

**"CGCAGACGTAGACGTCGTCGAGCAGCT"** was inserted onto GFP::GCN4-pLi before GCATcc (highlighted in bold)

4. Sequence of GFP::GCN4-pLi with positive charge:

**"CGCAGACGTAGACGTCGTCGAGCACGTCGTCGAGCACGTCGACGTGCCCGTGCTCGTCGTCGTCGAGCACGTCGTCGAGC
A"** was inserted onto GFP::GCN4-pLi before GCATcc (highlighted in bold)

5. Sequence of GFP::tba-2(T349E):
ATGCCTGCATccTCCAAGGGAGAGGAGCTCTTCACCGGAGTCGTCCCAATCCTCGTCGAGCTCGACGGAGgtattttcctgcatttttcaactg
ggaaaatgaaagaaaatcgataatttcagACGTCAACGGACACAAGTTCTCCGTCTCCGGAGAGGGAGAGGGAGACGCCACCTACGGAAAG
CTCACCCTCAAGTTCATCTGCACCACCGGAAAGCTCCCAGTCCCATGGCCAACCCTCGTCACCACCTTCTGCTACGGAGgtaagat
atgggaaagaaggaaaaaaccgagattttacttgaaaaattgaatttttcgcgggattttcaccaaaaattgttgaatattcattatttcacgctgtaaaacaaaaaaaaaaaaaatcaaaaa
ctacgttgaaatcgcgtttttaagcgaattttcttcagaattgccagattttaaccccaaattttgcagtttttaaataaaatttcaccttttcggctcaaattgtagattttcctgaaaatttagtacaaaa
aacaatttcctcgtaaatttttcaaattttcagTCCAATGCTTCTCCCGTTACCCAGACCACATGAAGCGTCACGACTTCTTCAAGTCCGCCATGCC
AGAGGGATACGTCCAAGAGCGTACCATCTTCTTCAAGGACGACGGAAACTACAAGACCCGTGCCGAGGTCAAGTTCGAGGGAG
ACACCCTCGTCAACCGTATCGAGCTCAAGGGAATCGACTTCAAGGAGGACGGAAACATCCTCGGACACAAGCTCGAGTACAACT
ACAACTCCCACAACGTCTACATCATGGCCGACAAGCAAAAGAACGGAATCAAGGTCAACTTCAAGgtacggattgaaattgctttaaaatttgaa
aaattgattaaaaagtgcatttttaagctttgaccgacttaaaattagatttctgagcctattttctgagaattggaattttttcattgtgaaagttcaaagaatagcgtgaatgattagaaaatattg
taaaatttcaatttttcctataaaaaaggatttttttaggaatcaaaaattgcaaaatgatgccctaaaattcgaaaataaatataaaaattggcggtttctcaaaaatctagaattccgacctta
attattaatttttttacaatattttttttgaaaaaaatccagaaaatttgaatttcgtagtttgtagtctaggcctctatcaattaaattttcgatttttgagtaaaatttcgaatttactattatttggaccaaa

attgtattttttttcagaattaaatttataaaattttcagaaattaaatttataaaaaaaaaaaaaaaattaaaaaaaaaaaaaaaaaattaaaaattaaaaattaaaaaattaagtttttttactca
aaattttgcactgaaattcgaaaatctaaaaatccgacctaaagtctgtatttttcaacaaaaattcagaaaaaactcaaaaacttgtattttgtagccagtcaaccactttctaaaatatcaaat
ttgaatttttcaagcatttttacattgaaaaatctaatttttcgagtgaaaccacttgaaaatctgaaaattgaattaatttctgaactttagagatttttgtcaaaattttagataaagggatttttaaac
aaaaaattgatttttaatcgaaaaatactgggatttatgggtttttaaaagaaaaacgggggtttgaatgaaaaatcgcctgaaatcttagaaaattaattaaaaaactatgatttaattccaa
aaattaccgaaaatatcaaattttccatttttaaaccctaaattctttcagATCCGTCACAACATCGAGGACGGATCCGTCCAACTCGCCGACCACTACCAA
CAAAACACCCCAATCGGAGACGGACCAGTCCTCCTCCCAGACAACCACTACCTCTCCACCCAATCCGCCCTCTCCAAGgtagatttttt
agaatttttgggtttttgaagtagaaaatcataaaaatctagggtttttatgaattgttttgaagaaaaattgcaaaaattccacaaaatggaagaaaaataactttggaagcgcatttttcgcaa
aaaaaccgaaatttttgcgtaaaatttcaaaattgcaataaaaattccacaaacatcaaatttcttaaattttttataaaaaattggatggaaacactcttgaatttagaaaaaaaatcagtttttct
catctaaaacttcaaaatttcggtgtaatcccattaaaattgccacaaaattcggaaatttcacctgaaatagagtgaataatttaaaatgttcagaaattcatattttgtcattttaaaagcattaa
aacaaatcaaaaatctattttttttggttggaaaagttcaaaattctggagaaattacattaaaattttcgtaaaatatggtaaataaacgaaaatgttgagaaattaaagaaaagttacaattttt
agcttaaaaattcaacattttgaggaaatgccacctaaaaaagtgactaatcgaaaatgttgagaaattaaaattgccaccatttatttatataaactactctaaaattacaattttcatgttaaaa
attaataaaaaatctactttttccaaactacagtaaccttaccgtatacctacagtacctgaacattgccccccaccagctcccaacccaatacctcctcaaaaacttacacctcaatttttcata
aactacagtaaccctaccaaaaaagcacaaaaaaaaatctacattcattttccaacaattttcaatattttcagGACCCAAACGAGAAGCGTGACCACATGGTCCTC
CTCGAGTTCGTCACCGCCGCCGGAATCACCCACGGAATGGACGAGCTCTACAAGCCTGCAATGCAATGCGTGAGGTCATCTCTA
TCCACGTCGGACAAGCCGGAGTCCAAATCGGAAACGCCTGCTGGGAGCTCTACTGCCTCGAGCACGGAATCCAGCCCGATGG
AACCATGCCAACTCAATCAACGAACGAGGGAGAGTCGTTCACCACTTTCTTCTCAGACACCGGATCCGGCCGTTACGTTCCAAG
ATCCATCTTCGTCGATCTCGAGCCAACTGTCGTTGACGAGATTCGCACTGGAACCTACAAGAAGCTCTTCCATCCAGAGCAGATG
ATCACCGGAAAGGAAGACGCCGCTAACAACTACGCTCGTGGACACTACACCGTCGGAAAGGAGCTCATCGACACCGTCCTCGA
CAGGATCCGTCGTCTCGCTGATAACTGCAGTGGACTCCAAGGATTCTTCGTCTTCCACTCCTTCGGAGGAGGTACCGGATCCGG
ATTCACTTCGCTTCTTATGGAACGTCTTTCCGTCGACTACGGAAAGAAGTCCAAGCTCGAGTTCTCCATCTACCCAGCTCCACAG
GTCTCAACCGCCGTCGTTGAGCCATACAACTCGATCCTCACCACCCATACCACCTTGGAGCACTCCGACTGCGCCTTCATGGTC
GATAACGAGGCCATCTACGACATCTGCCGCAGAAACTTGGATGTTGAGCGACCAAGCTACACCAACCTCAACAGAATCATCTCCC
AGgtttgttgagctcaatttgattgttgtattctaatgttctctttttacagGTTGTCTCCTCAATCACTGCTTCCTTGAGATTCGATGGAGCCCTCAACGTTGAT
CTCAACGAGTTCCAGACCAACTTGGTGCCATACCCAAGAATTCACTTCCCATTGGCCGCCTACACTCCACTCATCTCTGCTGAGA
AGGCCTACCACGAGGCTCTGTCCGTCAGCGACATCACCAATAGCTGCTTCGAGCCGGCTAACCAGATGGTCAAGTGTGATCCAC
GTCACGGAAAGTACATGGCTGTGTGCCTCTTGTACAGAGGAGACGTCGTTCCAAAGGACGTTAACACCGCCATCGCTGCAATCA
AGACCAAGAGAACCATCCAATTCGTCGATTGGTGCCCA**GAG**GGATTCAAGGTCGGAATCAACTACCAGCCACCAACTGTTGTGC
CAGGAGGTGATCTTGCCAAGGTGCCACGCGCCGTCTGCATGCTCTCCAACACTACCGCCATCGCTGAGGCCTGGTCTCGTCTC
GACTACAAGTTCGACTTGATGTACGCCAAGCGTGCCTTCGTCCACTGgtatgttgcccgttaaactatacttttcaaatattcaatgtgtttcttttcagGTACG
TCGGAGAAGGTATGGAGGAAGGAGAGTTCACCGAGGCTCGTGAGGACTTGGCTGCTCTCGAGAAGGACTACGAAGAGGTCGG
AGCTGACTCCAACGAGGGAGGAGAAGAGGAGGGAGAGGAGTACTAGCGCGTCGCGTAAaTAAaTAA

GFP optimized for germline expression is highlighted in green. tba-2(T349E) is colored red. T349E mutation is emphasized in bold text.

Table S1: *C. elegans* strains used in this study.

| Strain name | Genotype | Source |
|---|---|---|
| FM917 | *fxIs1 [pie-1p::TIR1::mRuby, I:2851009] I;<br>ltIs37 [pAA64; pie-1p::mCh::his-58 + unc-119(+)];<br>ruIs57 [pie-1p::GFP::tubulin + unc-119(+)] V* | This study |
| FM1054 | *fxIs1[pie-1p::TIR1::mRuby, I:2851009] I;<br>ran-3(syb7781[ ran-3-3xGAS-AID-3xGAS-HALO]) II;<br>ltIs37 [pAA64; pie-1p::mCh::his-58 + unc-119(+)];<br>ruIs57 [pie-1p::GFP::tubulin + unc-119(+)] V* | This study |
| FM1056 | *fxIs1[pie-1p::TIR1::mRuby, I:2851009] I;<br>ran-2(syb7819[ran-2-3xGAS-AID-3xGAS-HALO]) III;<br>ltIs37 [pAA64; pie-1p::mCh::his-58 + unc-119(+)];<br>ruIs57 [pie-1p::GFP::tubulin + unc-119(+)] V* | This study |
| FM717 | *bus-17(e2800)X;<br>ltSi1412 [pNA20; Pmex-5::mNeonGreen::tbb-2 operon linker mCh::his-11::Ptbb-2; cb-unc-199(+)]I;<br>unc-119(ed3)III clone B (MOS I insertion)* | This study |
| NM5402 | *jsSi1579 [loxP::rpl-28p::FRT::GFP::his-58 FRT3] II.<br>bqSi711 [mex-5p::FLP::SL2::mNG + unc-119(+)] IV.* | Gifted from Nonet Lab |
| FM971 | *duSi18[GFP(SMU)-GCN4-pLI] II;<br>ltIs37 [pAA64; pie-1p::mCh::his-58 + unc-119(+)];<br>him-8(e1489)* | This study |
| FM1011 | *duSi20[GFP(SMU)::tba-2(T349E)] II<br>ltIs37 [pAA64; pie-1p::mCh::his-58 + unc-119(+)];<br>him-8(e1489)* | This study |
| FM628 | *unc-119(ed3) III;<br>ltSi464[pNH103; Pmex-5::npp6::GFP::tbb-2 3'UTR; cbunc-119(+)] I;<br>ltIs37[pAA64; pie-1::mCherry::his-58; unc-119 (+)] IV* | A gift from Oegema-Desai Lab |
| BN359 | *ima-2(ok256) I/hT2[bli-4(e937) let-?(q782) qIs48] (I;III);<br>qaIs3502[pie-1p::YFP::lmn-1 + pie-1p::CFP::H2B + unc-119(+)]* | CGC |
| FM991 | *wjIs76[Cn_unc-119(+); pie-1p::mKate2::tba-2];<br>vit-2(crg9070[vit-2::gfp]) X;<br>egxSi126 [mex-5p::hsp-3(aa1-19)::halotag::HDEL::pie-1 3'UTR+ unc-119(+)] I. "* | This study |
| FM691 | *cox-4(zu476[cox-4::eGFP::3xFLAG]) I;<br>wjIs76[Cn_unc-119(+); pie-1p::mKate2::tba-2]* | This study |
| CZ18550 | *juSi123[rpl-29::GFP] II; rpl-29(tm3555) IV* | CGC |
| ABR5 | *staIs1 [pie-1p::GFP + unc-119(+)];<br>unc-119(ed3) III* | CGC |

| | | |
|---|---|---|
| FM1103 | *duSi21[HALO(smu)] II;*<br>*ruIs57 [pie-1p::GFP::tubulin + unc-119(+)] V*<br>*itIs37 [pie-1p::mCh::H2B::pie-1 3'UTR + unc-119(+)] IV"* | This study |
| FM1168 | *duSi23[minus7-GFP(SMU)::GCn4-pLI] II;*<br>*ItIs37[pAA64; pie-1::mCherry::his-58; unc-119 (+)] IV* | Negative Charge; this study |
| FM1169 | *duSi24[plus7-GFP(SMU)::GCn4-pLI] II;*<br>*ItIs37[pAA64; pie-1::mCherry::his-58; unc-119 (+)] IV* | Neutral Charge; this study |
| FM1180 | *duSi25[plus21-GFP(SMU)::GCn4-pLI] II;*<br>*ItIs37[pAA64; pie-1::mCherry::his-58; unc-119 (+)] IV* | Positive charge; this study |
