## Supplementary figures and images for "Roles of Tubulin Concentration during Prometaphase and Ran-GTP during Anaphase of *C. elegans* meiosis"

### Figure S1

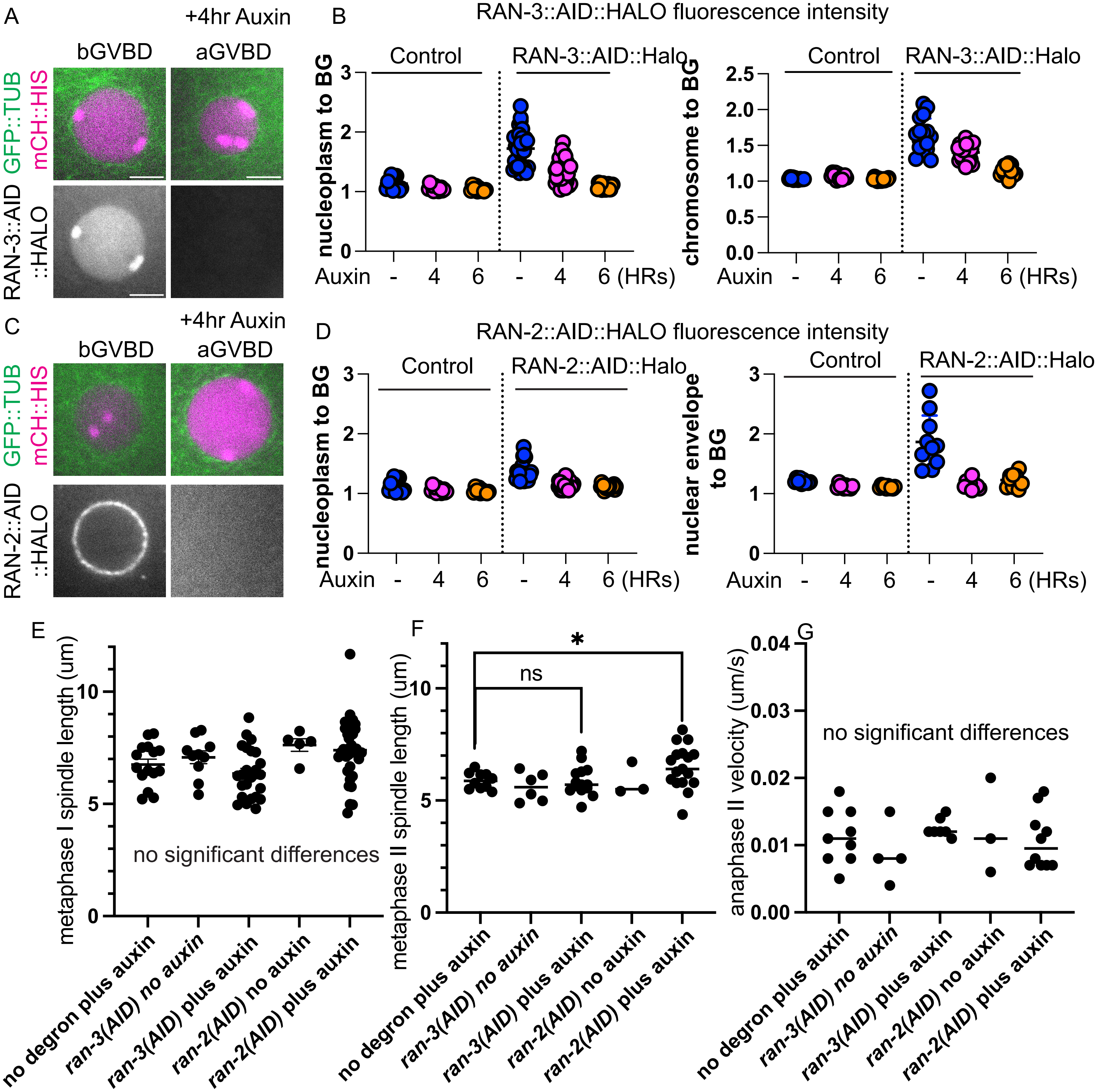

### Figure S2

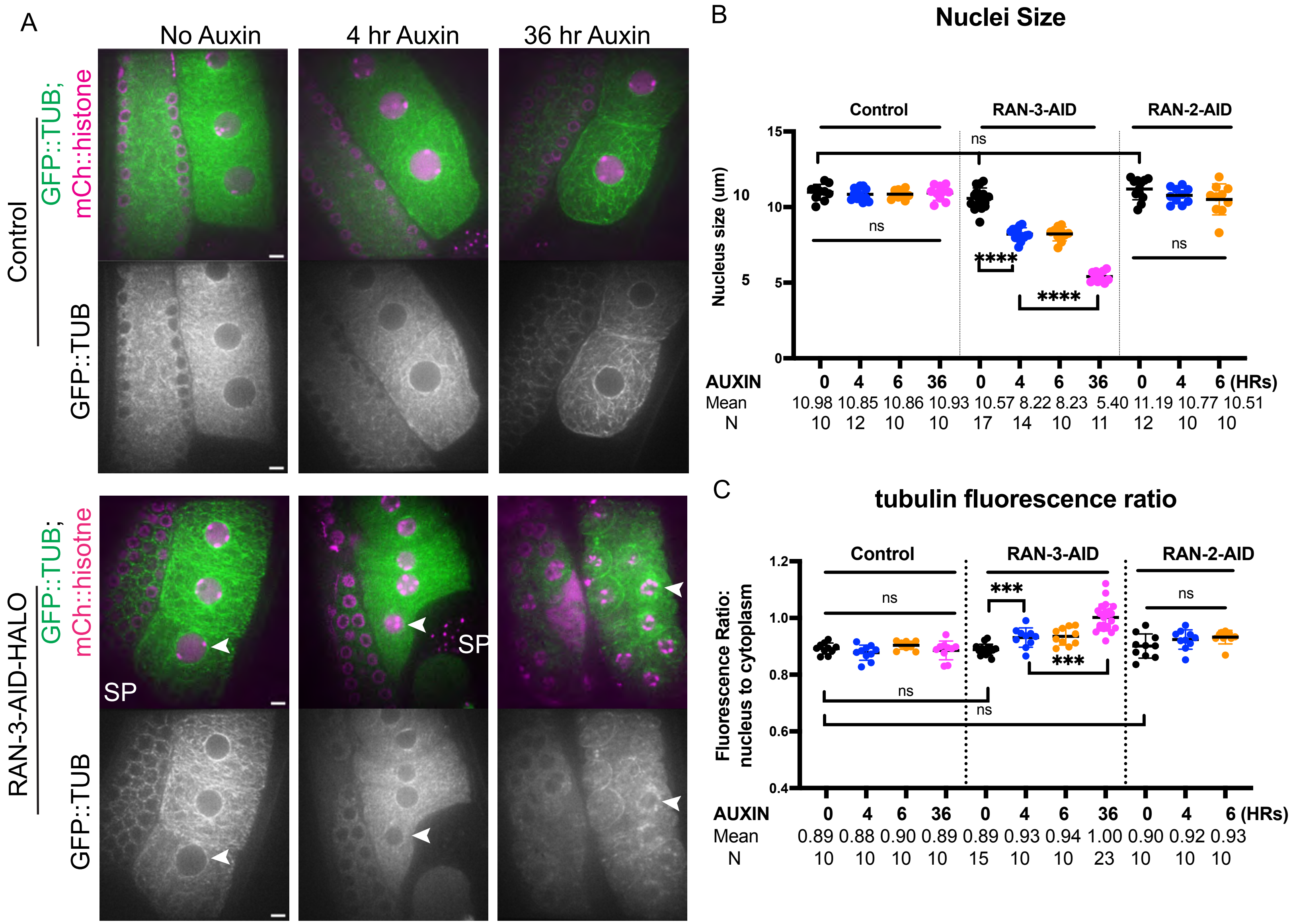

### Figure S3

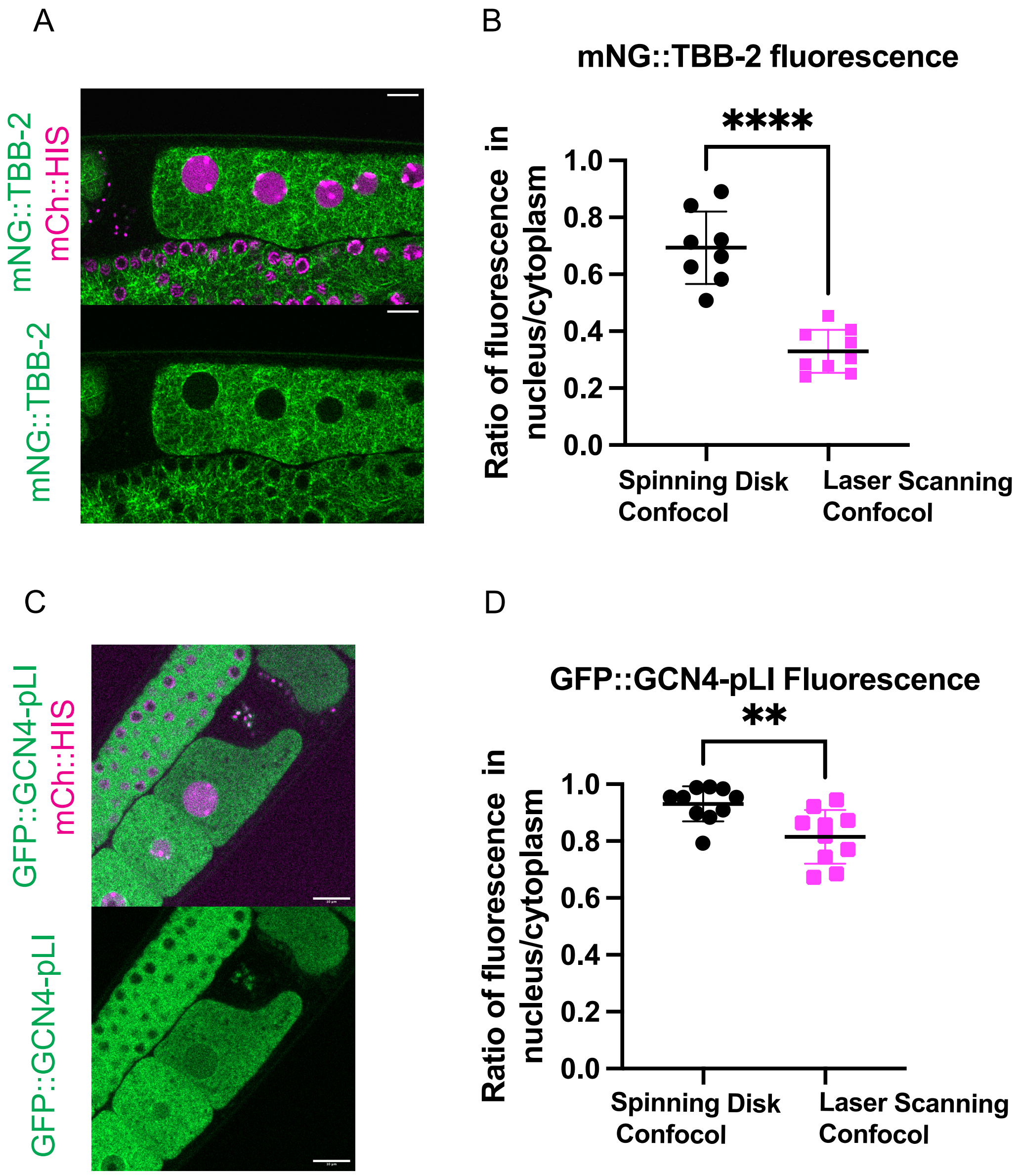

### Figure S4

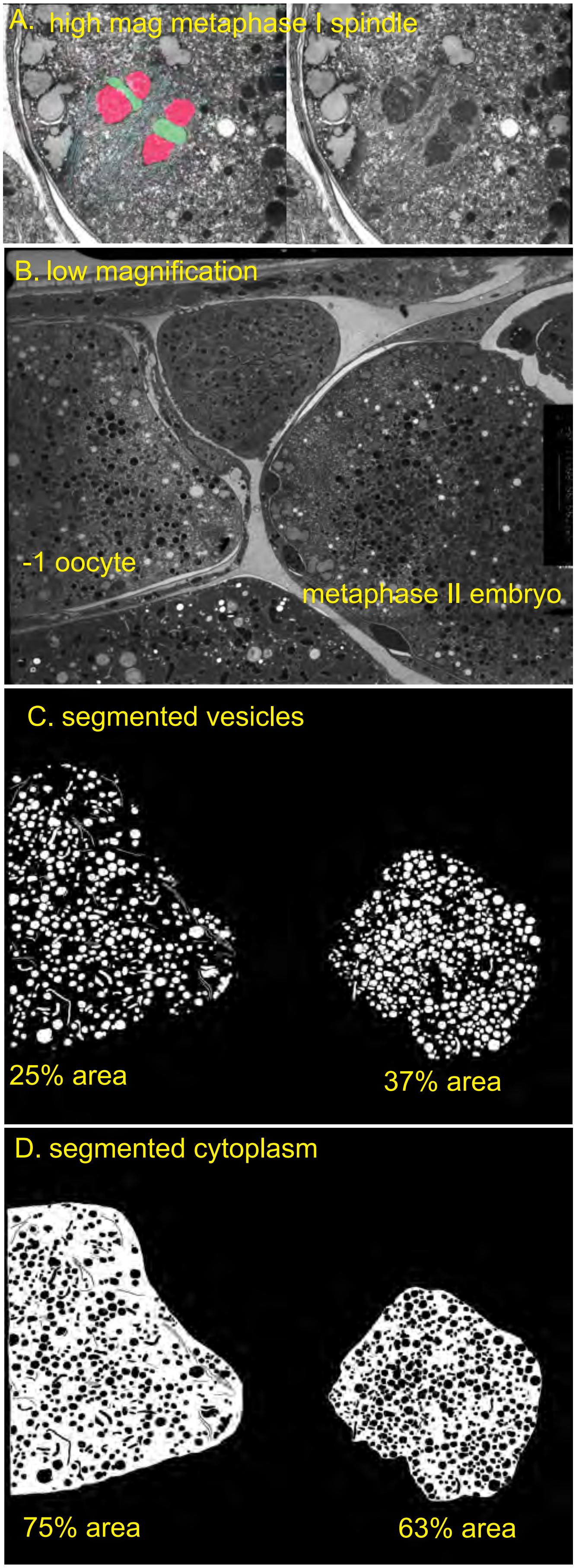

### Figure S5

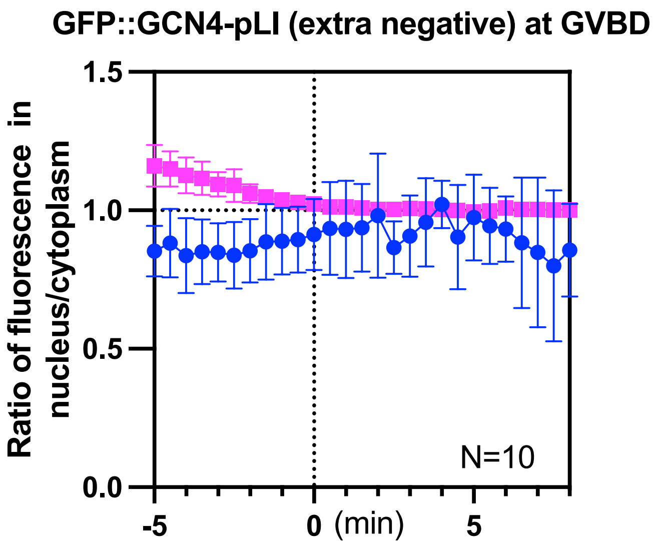
